# Macrophages provide a transient muscle stem cell niche via NAMPT secretion

**DOI:** 10.1101/2020.08.24.262428

**Authors:** Dhanushika Ratnayake, Phong D. Nguyen, Fernando J. Rossello, Verena C. Wimmer, Abdulsalam I. Isiaku, Laura A. Galvis, Alasdair J. Wood, Ziad Julier, Thomas Boudier, Viola Oorschot, Kelly L. Rogers, Mikaël M. Martino, Christophe Marcelle, Graham J. Lieschke, Jeroen Bakkers, Peter D. Currie

## Abstract

Skeletal muscle is paradigmatic of a regenerative tissue that repairs itself via the activation of a resident stem cell^1^. Termed the satellite cell, these normally quiescent cells are induced to proliferate by ill-defined wound-derived signals^2^. Identifying the source and nature of these pro-regenerative cues has been hampered by an inability to visualise the complex cellular interactions that occur within the wound environment. We therefore developed a zebrafish muscle injury model to systematically capture satellite cell interactions within the injury site, in real time, throughout the repair process. This analysis identified that a specific subset of macrophages ‘dwells’ within the injury, establishing a transient but obligate stem cell niche required for stem cell proliferation. Single cell profiling identified specific signals secreted from dwelling macrophages that include the cytokine, Nicotinamide phosphoribosyltransferase (NAMPT/Visfatin/PBEF). Here we show that NAMPT secretion from the macrophage niche is required for muscle regeneration, acting through the C-C motif chemokine receptor type 5 (CCR5) expressed on muscle stem cells. This analysis reveals that along with their well-described ability to modulate the pro-inflammatory and anti-inflammatory phases of wound repair, specific macrophage populations also provide a transient stem cell-activating niche, directly supplying pro-proliferative cues that govern the timing and rate of muscle stem cell-mediated repair processes.

## Main

Live imaging of the collective cellular response to tissue injury remains a long-term goal of the regenerative medicine field. In an attempt to attain this goal we have developed zebrafish models of tissue injury where the optical accessibility of the larvae allows the application of non-invasive techniques to assay repair in real time. Here we apply this approach to regenerating skeletal muscle in order to determine the cellular and molecular events that define regeneration *in vivo*. Transgenic zebrafish reporter lines fluorescently tagging wound-present cellular components were subject to acute injury, enabling the location and response dynamics of individual wound-occupying cells to be correlated to the stem cell compartment during muscle repair. We developed two injury paradigms: a focal laser ablation model that severs 2-6 fibres per injury and a needle stab model that consistently injures 2 adjacent myotomes. Systematic characterisation of different cellular behaviours within the regenerating niche in both models highlighted the directed and obligate association between specific macrophage populations and muscle stem cells during repair. Distinct modes of high-resolution imaging coupled with *in vivo* cell tracking defined these interactions with high temporal and spatial resolution.

Initially, multiphoton imaging was utilised to capture the response prior to, and immediately following, laser ablation muscle injury. The compound transgenic line Tg(*actc1b*:GFP); Tg(*mpeg1*:GAL4FF/*UAS*:NfsB-mCherry), in which differentiated muscle expresses GFP and macrophages are labelled by mCherry fluorescence, allowed the wound region to be defined while observing the kinetics of injury-responsive macrophages (Fig. 1a-c, Supplementary video. 1). Following muscle fibre laser ablation injury, 34±2% of macrophages (MΦs) in the vicinity of the wound (MΦs within a 260 μm radius from the centre of the laser ablation, a region encompassing 2 myotomes on either side of the injury, n=4 injuries) displayed rapid, active and directed migration towards the wound site (average distance travelled: 128.31±68.03 μm, average velocity: 0.149±0.040 μmsec^-1^, n=30 MΦs assayed in n=4 larvae, Fig. 1c, Supplementary Video. 1). That not all the injury-proximate macrophages responded to the lesion suggested the potential existence of specific macrophage subsets that are primed to respond to trauma.

**Figure 1.**
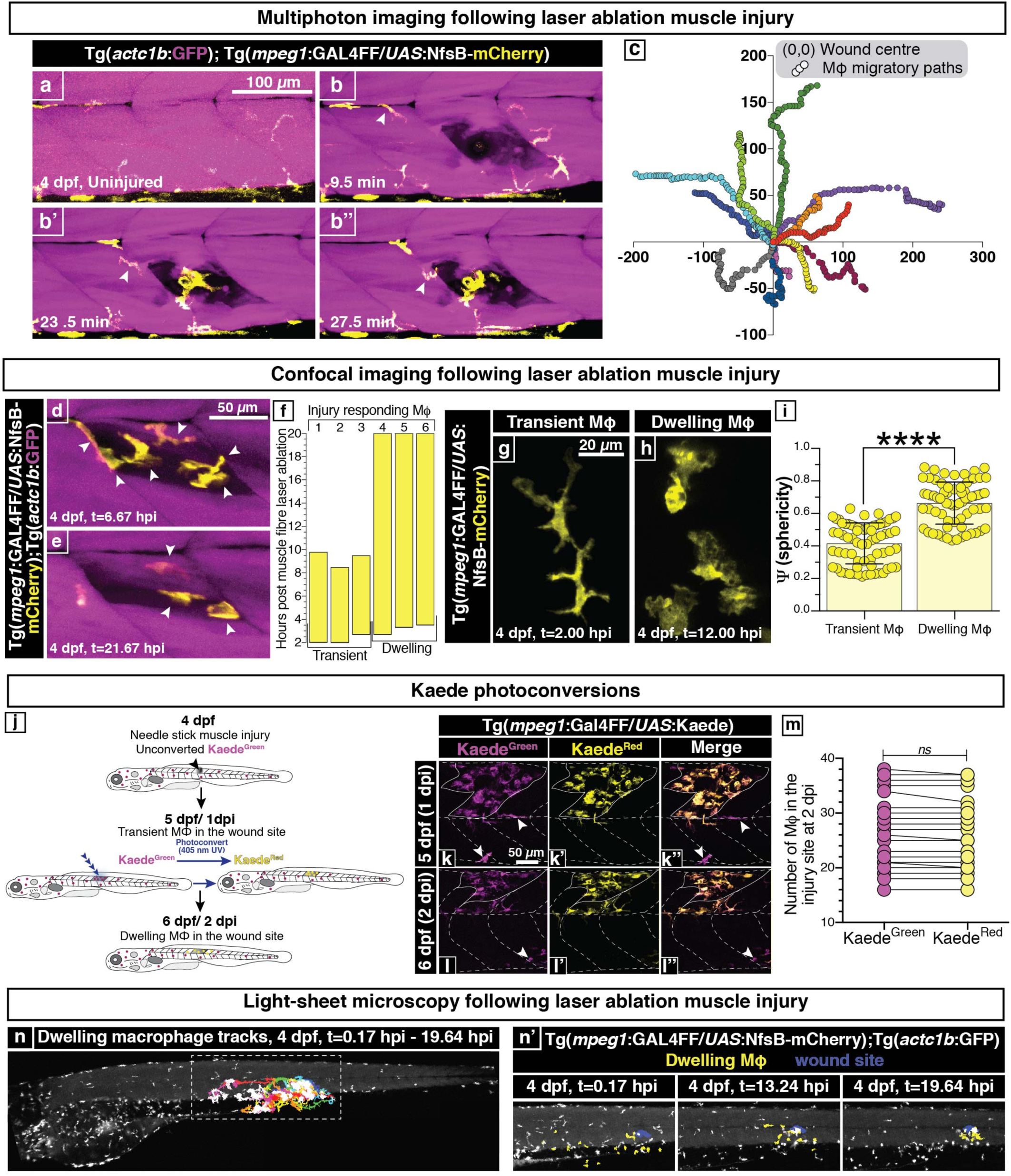
A subset of injury-responsive macrophages dwell at the wound site for the duration of repair. **a-c**, Uninjured muscle (Tg(*actc1b*:GFP), magenta) is patrolled by macrophages (MΦs) (Tg(*mpeg1*:GAL4FF/*UAS*:NfsB-mCherry), yellow) at 4 dpf (**a**). Upon injury, MΦs are activated and rapidly migrate into the wound site (**b-b’’**, arrowheads). Frames extracted from Supplementary Video. 1 (n=3). (**c**) The migratory paths of injury-responding MΦs are graphed, where (0,0) is set to the centre of the wound. **d-f**, *In toto* imaging identifies two MΦ subsets occupying the wound at distinct temporal phases (**f**), an early injury-responsive transient (**d**) and late injury-located dwelling subset (**e**) (arrowheads). Frames extracted from Supplementary Video. 2 (n=8). **g-i**, Morphological characterisation of these two subsets revealed transient MΦs are more stellate (**g**) with lower sphericity (**i**), while dwelling MΦs are more rounded (**h**) with higher sphericity (**i**) (n=69 MΦs per group, extracted from n=18 Videos). Mean±S.D. Significance (\*\*\*\**P*<0.0001) in unpaired *t*-test. **j-m**, Tracking of wound located MΦs reveals that dwelling MΦs are derived from transient MΦs. (**j**) Photoconversion strategy schematic. At 1dpi, injury-located MΦs in Tg(*mpeg1*:Gal4FF/*UAS*:Kaede) larvae were photoconverted (**k-k’’**, pre conversion; magenta (arrowheads), post conversion; yellow), reassessed at 2 dpi (**l-l’’**) and quantified (**m**, n=20). Not significant (ns) in unpaired *t-*test. **n-n’**, Retrospective tracking of injury-present dwelling MΦs at 24 hpi identified their injury-proximate origin. (**n**) Dwelling MΦ track projections for the duration of imaging are superimposed on whole larvae. (**n’**) The location of dwelling MΦs (yellow) at different time points following imaging demonstrates their origin and sequential migration into the injury site (blue). Frames extracted from Supplementary Video. 4 (n=4).

Next, injury-migrated macrophage dynamics within the boundaries of the wound site were examined at a cellular resolution following muscle fibre laser ablation. High-resolution continuous confocal imaging demonstrated that peak macrophage numbers within the wound site were reached at 2.50±0.42 hours post injury (hpi) (n=8 injuries), following which, no additional macrophages migrated into the wound (Fig. 1d-f, Supplementary Video. 2). Of the total injury-responding macrophages, 51.11±1.83% (n=8 injuries) remained within the wound site up to 24 hpi (Fig. 1e-f). We refer to these long-term injury-located macrophages as ‘dwelling’. In contrast, macrophages that exit the wound site do so prior to 10.48±1.19 hpi and were consequently designated ‘transient’ (Fig. 1d, f). High-resolution confocal imaging revealed that these two macrophage populations display morphological differences, with the transient cells exhibiting an obvious stellate appearance while the dwelling cells adopt a more spherical form (dwelling MΦs exhibited a 0.248±0.022 higher sphericity value when compared to transient MΦs, Fig 1g-i, Supplementary Video. 3). This transient to dwelling macrophage transition occurred irrespective of the magnitude of injury, but scaled temporally with wound size (51.6% of needle stab injury-responsive MΦs went on to dwell by 2 days post injury (dpi), n=20, Extended Data Fig. 1a-d).

To establish whether dwelling macrophages are a subset of the original pool of injury-responsive macrophages, a Tg(*mpeg1*:Gal4FF/*UAS*:Kaede) transgenic line in which macrophages are labelled with the photoactivatable fluorescent protein Kaede was utilised. One-day post needle stick muscle injury, transient macrophages present within the wound site were photoconverted to distinguish them from macrophages external to the injury (n=20, Fig. 1j-k’’). On re-imaging the injury site one day later, all the dwelling macrophages within the wound site exhibited the photoconverted form of Kaede (Fig. 1l-m). This indicates that dwelling macrophages are a subset derived from the initial transient injury-responding macrophage population, and not the result of another, temporally distinct migration.

Further investigations into the nature of the macrophages that become injury dwelling were carried out in order to distinguish their origin as either tissue resident and/or migratory. To address this question, macrophages throughout the entire larvae were tracked from 10 min to 25 hpi following laser ablation muscle fibre injury in the Tg(*actc1b*:GFP) and Tg(*mpeg1*:GAL4FF/*UAS*:NfsB-mCherry) transgenic line by whole larval light-sheet microscopy (Supplementary Video. 4). Retrospective tracking of injury-located macrophages revealed that all macrophages that proceeded to assume a dwelling phenotype migrated from the vicinity of the wound site, and as such are tissue resident rather than migratory in nature (n=4 injuries, Fig. 1n-n’, Supplementary Video. 4). These observations suggest that there is regionalised anatomical constraint to the activation of wound-responsive macrophages in the zebrafish larvae. However, why some wound-proximate macrophages refrain from migrating into the injury site remains an open question.

As the residency of dwelling macrophages occurred in the same temporal window as stem cell mediated muscle repair^3^, we examined if they were required for this process by comparing muscle regeneration in the presence and absence of the transient and dwelling macrophage populations. The Tg(*mpeg1*:GAL4FF/*UAS*:NfsB-mCherry) transgenic line allows temporally-controlled nitroreductase-mediated genetic ablation of macrophages by the addition of metronidazole (Mtz) at discrete time points during muscle regeneration (Fig. 2a, Extended Data Fig. 2a-c). Regeneration following needle stick muscle injury was assessed by birefringence imaging of the wound site at key milestones over the course of the repair process (n=24 per group, Fig. 2b-h). The pseudo-crystalline array of muscle sarcomeres confers an intrinsic birefringence, making uninjured muscle appear bright against a black background when observed using polarised light, enabling non-invasive visualisation of muscle integrity^4^. Addition of Mtz at the time of injury (to ablate all injury responding macrophages) and Mtz treatment at 1.75 dpi (to preferentially ablate dwelling macrophages) both resulted in a significant regeneration deficit (Fig. 2b-h). Hence, not only are macrophages necessary for efficient skeletal muscle regeneration, but there is also an explicit requirement for the dwelling macrophage subpopulation for injury repair (Extended Data Fig. 2d-g).

**Figure 2.**
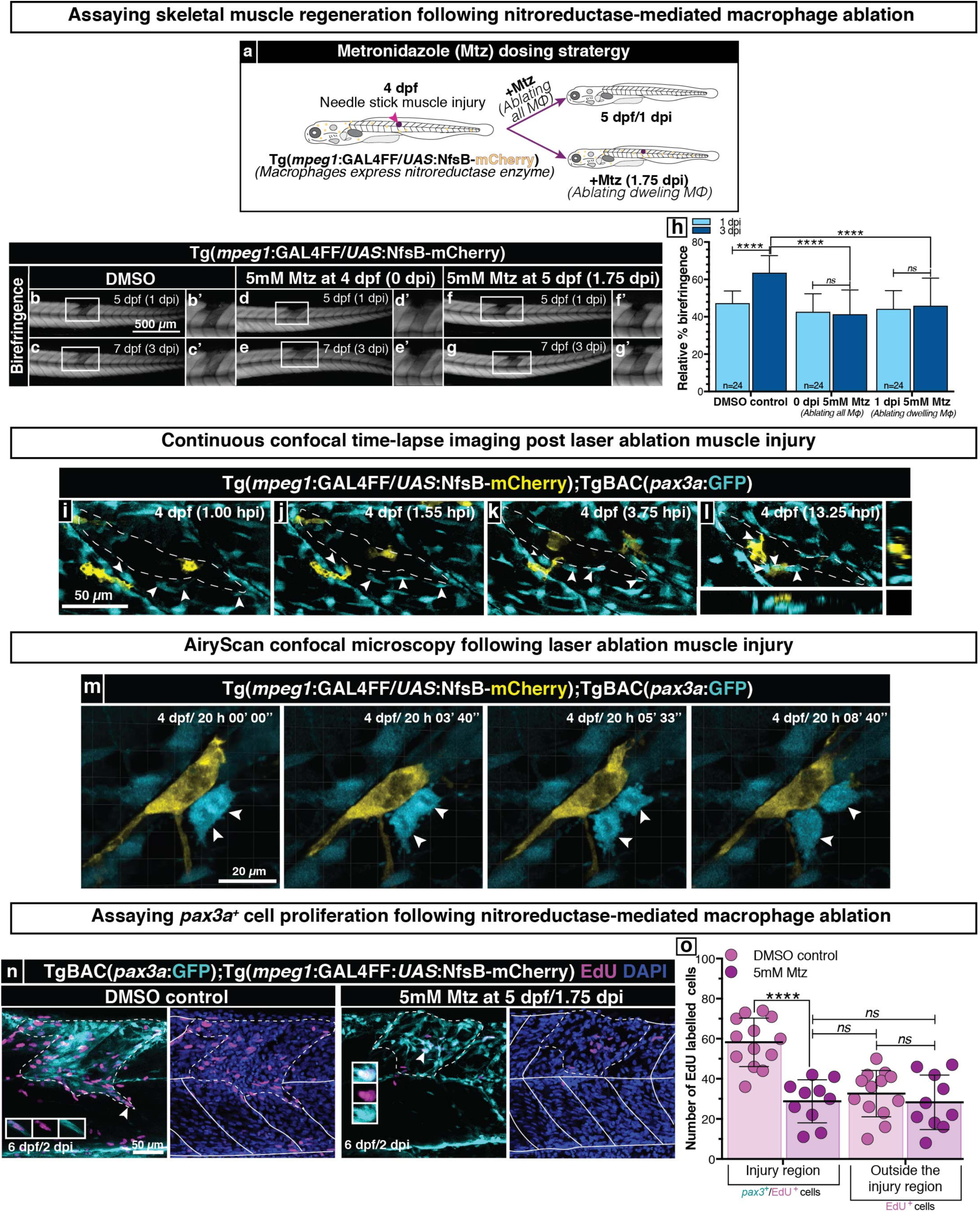
Dwelling macrophages induce muscle stem cell proliferation *in vivo*. **a**, Dosing strategy schematic for addition of the pro-drug metronidazole (Mtz) to ablate either all MΦs, or specifically dwelling MΦs in Tg(*mpeg1*:GAL4FF/*UAS*:NfsB-mCherry) larvae. **b-h**, Both these ablation strategies result in a significant regeneration deficit as observed by birefringence imaging (**b-g’**) and quantification (**h**, n=24). **i-l**, Following laser ablation injury (dotted line) transient MΦs migrate into the wound site (Tg(*mpeg1*:GAL4FF/*UAS*:NfsB-mCherry), yellow) (**i-k**). Simultaneously, activated *pax3a*^+^ myogenic stem cells (TgBAC(*pax3a*:GFP), cyan) from the injured and adjacent myotomes travel to line the edge of the injury site (**i-k**, arrowheads). Once MΦs transition to a dwelling subtype they initiate interactions with wound edge lining *pax3a*^+^ cells (**l**, orthogonal views). Frames extracted from Supplementary Video. 5 (n=5). **m**, High-resolution AiryScan microscopy of these interactions reveal dwelling MΦs maintain prolonged intimate associations with *pax3a*^+^ muscle stem cells (arrowheads), following which the associated stem cell undergoes cell division. Frames extracted from Supplementary Video. 6 (n=10). **n-o**, EdU incorporation assessment of stem cell proliferation following dwelling MΦ ablation reveals a requirement for this MΦ subset to maintain muscle stem cell divisions in the wound site. The homeostatic level of myotome cell proliferation external to the injury zone provides an internal specificity control for these analyses, with no significant difference in cell proliferation in the presence or absence of dwelling MΦs being observed outside the wound. Quantification (**o**). Mean±S.D. Significance (\*\*\*\**P*<0.0001) in two-way ANOVA with Tukey’s multiple comparison test.

Wound debris clearance is mediated by a number of cell types^5^, but largely attributed to phagocytic macrophages^6^. Absent or ineffective clearance might be one explanation for the lack of regeneration observed when macrophage populations are ablated early in the regenerative process. Three separate approaches were undertaken to assay wound debris clearance when all injury responding macrophages were ablated. Firstly, larvae were stained with fluorescently conjugated phalloidin to detect the actin cytoskeleton of remnant necrotic skeletal muscle fibres in the injury site. Secondly, LysoTracker fluorescent vital dye was used to visualise acidic membranous organelles and the autophagic process and finally, the Tg(*ubi*:secAnnexinV-mVenus) transgenic line was utilised to identify apoptotic cells within the injury zone. In none of these independent approaches did we observe accumulation of cellular debris in the injury site of macrophage-ablated larvae compared to un-ablated controls (Extended Data Fig. 3a-e). These data suggest that the failure of repair evident in macrophage-deficient settings does not result from ineffective debris clearance, but rather via the lack of an as-yet uncharacterised pro-myogenic function.

**Figure 3.**
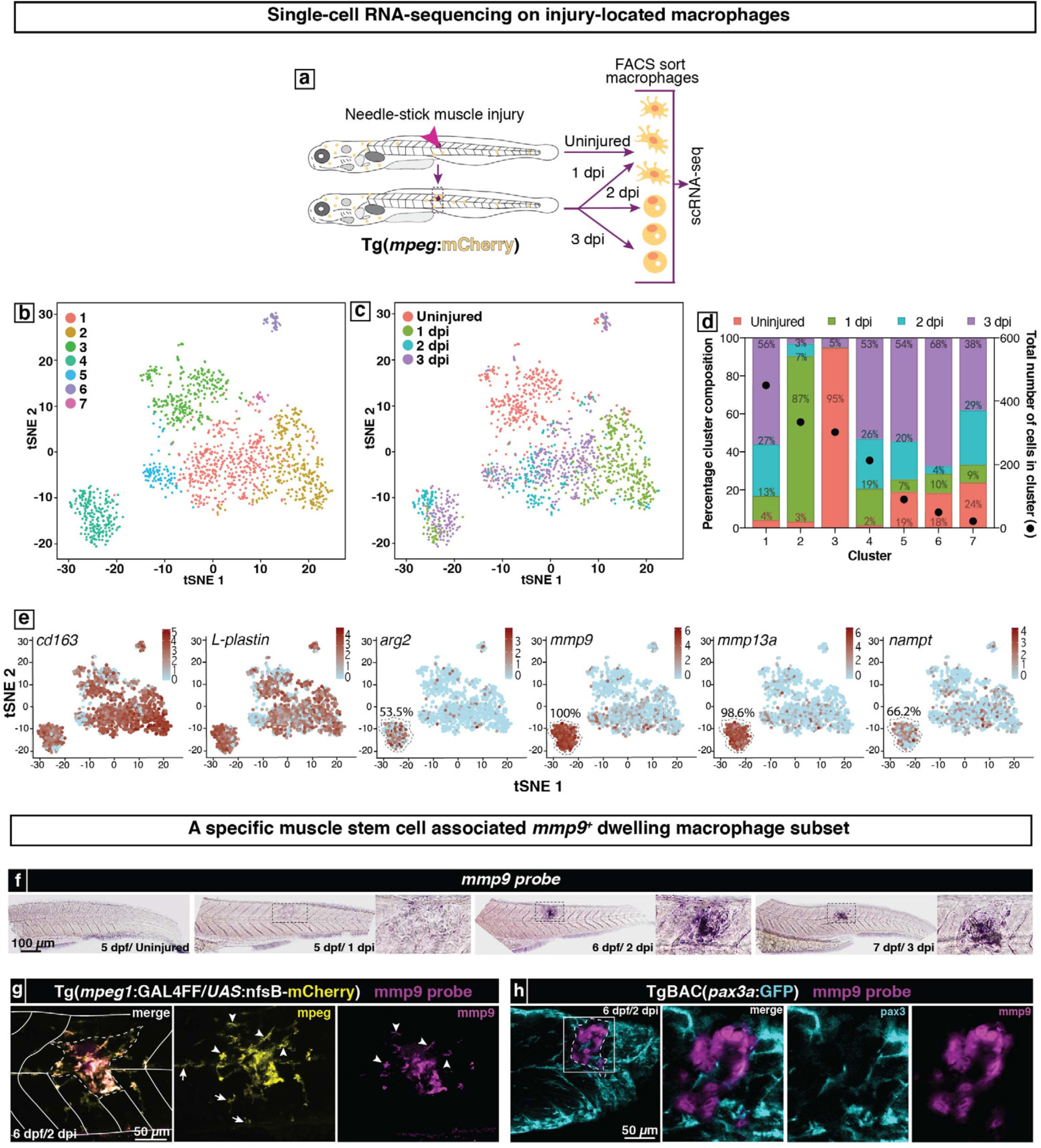
Single-cell RNA-seq identifies a unique *mmp9* positive dwelling macrophage subset. **a**, Schematic of injury-responsive macrophage scRNA-seq workflow. **b**, tSNE plot revealing spontaneous cell clusters. **c**, In this graph the injury time point of isolated MΦs is overlaid on the tSNE plot and illustrates the correspondence between unsupervised clustering and the described transient and dwelling MΦ subtypes. **d**, Uninjured MΦs cluster together (Cluster 3). Macrophages isolated from a ‘transient’ time-point (1dpi) also predominantly cluster together (Cluster 2). The remaining 5 clusters (1, 4, 5, 6, 7) are predominated by macrophages isolated from ‘dwelling’ time points (2-3 dpi) and display a surprising degree of molecular heterogeneity. **e**, The macrophage identity of sorted cells was validated by their collective expression of the pan-macrophage markers *cd163* and *L-plastin*. Known pro-regenerative MΦ markers *arg2, mmp9* and *mmp13a* are concentrated in cluster 4 present MΦs. The percentage of cells in Cluster 4 expressing these markers is shown. In addition, this cluster also differentially expresses *nampt. G*ene expression levels are log-normalised gene read counts using a scaling factor of 10,000. **f**, The spatiotemporal expression of *mmp9* RNA was assayed by *in situ* hybridisation following needle stick injury. *mmp9* expression is selectively up-regulated in the wound site from 2 dpi onwards (n>15). **g-h**, *mmp9* expression (magenta) in the wound site was spatially resolved and identified to be present in a subset of dwelling MΦs (yellow, arrowheads, MΦs not expressing *mmp9*; arrows, wound site: dotted line, n=15) (**g**) that are associated with wound present *pax3a*^+^ stem cells (cyan, wound site: dotted line, n=15) (**h**).

Given that the position that dwelling macrophages occupy within the repair niche was topographically similar to that which we had previously identified as containing dividing muscle stem and progenitors cells^3^ (Fig. 1e), we examined the spatial and temporal relationship between these two populations of cells during muscle regeneration. Long-term confocal time-lapse imaging following laser ablation muscle injury was performed in the compound Tg(*mpeg1*:GAL4FF/*UAS*:NfsB-mCherry);TgBAC(*pax3a*:GFP) transgenic line which labels both macrophages and *pax3a* expressing muscle stem/progenitor cells (Fig. 2i-l, Supplementary Video. 5). The *pax3a*:GFP reporter is particularly useful in this context as it is expressed in both the muscle stem cell compartment as well as the dividing progenitor population^7^. Following injury, transient macrophages and *pax3a*^+^ cells both responded and migrated simultaneously to the wound. However, *pax3a*^+^ cells migrated independently of transient macrophages, displaying distinct migration kinetics, taking residence in their niche at the edge of the wound by 10.25±1.99 hpi (n=5 injuries, Fig. 2i-l, Extended Data Fig. 4a-b’, Supplementary Video. 5). Following their transition from a transient to dwelling macrophage phenotype, dwelling macrophages associated with the *pax3a*^+^ cells lining the wound edge at 11.17±1.13 hpi and displayed continuous interactions with adjacent stem cells over the course of 5.38±1.79 h (Fig. 2l, Supplementary Video. 5).

**Figure 4.**
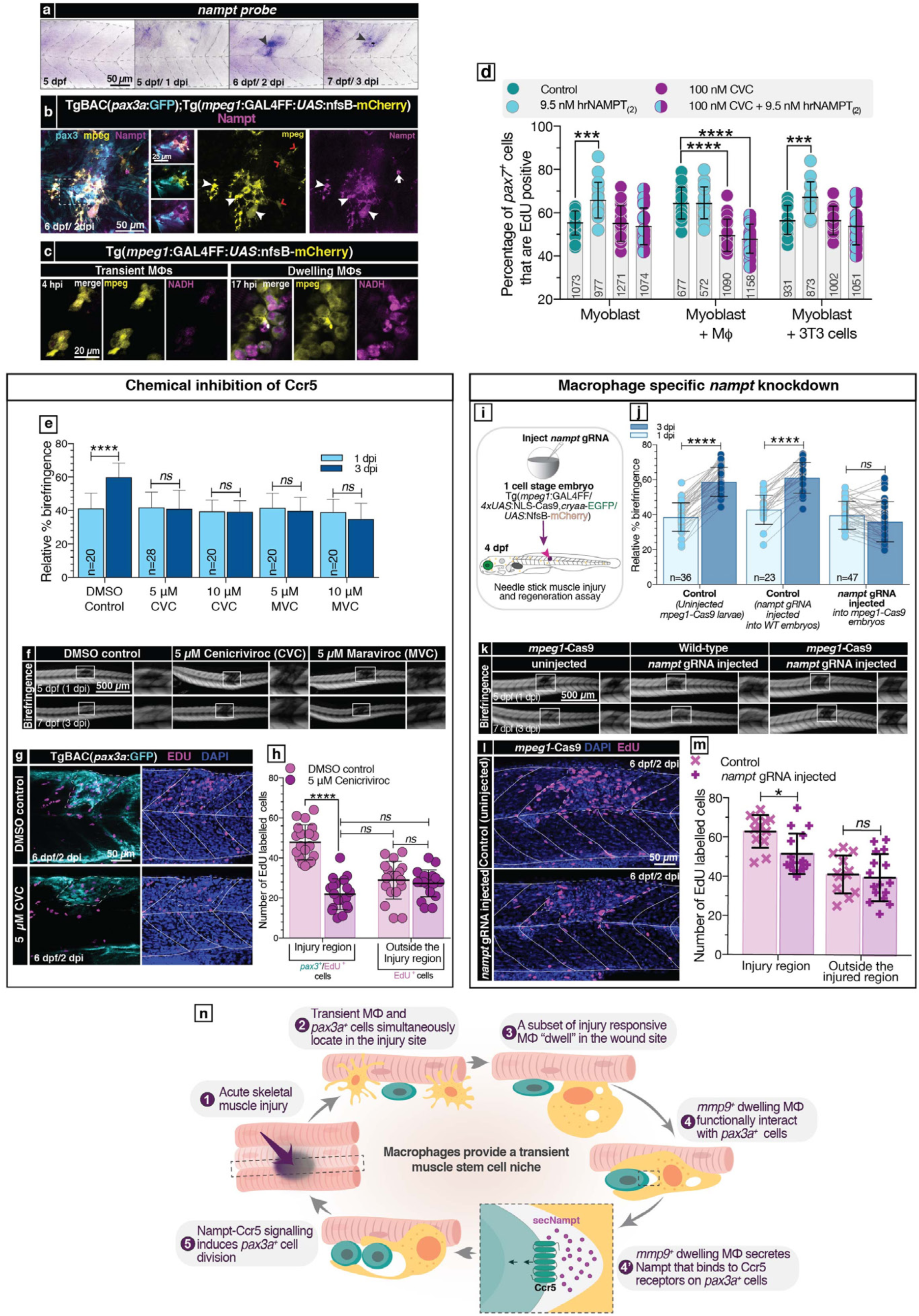
Macrophage secreted Nampt binding to Ccr5 induces stem cell proliferation. **a**, *nampt* mRNA expression is specifically up-regulated in the injury site from 2 dpi onwards (arrowhead, n>15). **b**, Within the injury site Nampt (magenta) is expressed by macrophages (yellow) associated with muscle stem cells (cyan) as well as being present in the extracellular space (n=15). **c**, NADH auto-fluorescence (magenta) displays a localised up-regulation in dwelling macrophages (yellow) indicating these cells as the primary source of wound present Nampt (n=15). **d**, NAMPT stimulates proliferation of PAX7^+^ satellite cells *in vitro*. Satellite cells in mouse primary myoblast monocultures display enhanced proliferation upon exogenous NAMPT supplementation. This pro-proliferative response is abolished upon administration of the CCR2/CCR5 dual inhibitor, cenicriviroc (CVC). Co-culturing myoblasts with macrophages stimulates satellite cell proliferation, a response that is inhibited upon CVC administration. Co-culturing myoblasts with 3T3 cells that do not secrete NAMPT does not stimulate satellite cell proliferation. Total number of assayed PAX7^+^ cells recorded in each bar of the graph. **e-f**, Similarly, larvae soaked in CVC or the Ccr5 specific inhibitor maraviroc (MVC) displayed a marked regeneration deficit revealed by birefringence imaging (**f**) and quantified (**e**, n=20). **g-h**, *pax3a*^+^ myogenic stem cell proliferation is inhibited by CVC addition as demonstrated by decreased EdU incorporation of these cells in the wound site post-injury (**g**) and quantified (**h**). **i-m**, Nampt’s specific role in injury-responsive macrophages was interrogated by means of a novel tissue specific gene knockout strategy (schematic, **i**). Macrophage-specific Nampt knockout larvae displayed a marked regeneration deficit (imaged (**k**) and quantified (**j**)) due to an inability to maintain the required proliferative response within the injury zone (imaged (**l**) and quantified (**m**)). **d**,**e**,**h**,**j**,**m**, Mean±S.D. Significance (\*\*\*\**P*<0.0001, \*\*\**P*<0.001, \*\**P*<0.01, \**P*<0.1) in two-way ANOVA with Tukey’s multiple comparison test. **n**, Schematic of injury-responding MΦs role in modulating muscle stem cell proliferation.

In order to better visualise these cellular interactions, high-resolution live imaging using AiryScan confocal microscopy was carried out on co-located dwelling macrophages and *pax3a*^+^ muscle stem/progenitor cells (Fig. 2m, Supplementary Video. 6). We observed sustained associations between the co-resident macrophages and stem cells. Macrophages maintained highly dynamic membrane contacts with the stem cells, enveloping them with continuous and repetitive membrane extensions (n=10 injuries, Fig. 2m, Extended Data Fig. 4c, Supplementary Video 6, 7). These macrophage-stem cell interactions were further validated by correlative light and electron microscopy (CLEM), which confirmed that the two cell types displayed tight membrane appositions in three-dimensional XYZ planes (Extended Data Fig. 5a-d). Following these protracted physical interactions, the associated *pax3a*^+^ muscle stem cell invariably underwent cell division (Fig. 2m, Extended Data Fig. 4c, Supplementary Video 6, 7). On average, macrophages interacted with a specific stem/progenitor cell for 5.42±1.72 h prior to the associated stem cell undergoing division. Crucially, no muscle stem cell divisions were observed in the absence of prolonged interactions with co-resident dwelling macrophages (n=26 wound associated stem cell divisions imaged in n=10 independent larvae). These data suggested that macrophage-stem cell associations maybe essential for stem cell proliferation.

To assay the requirement of dwelling macrophages to stimulate stem cell proliferation we performed EdU labelling in myotome needle stick injured larvae where dwelling macrophages had been ablated (Fig. 2n-o). Ablation of dwelling macrophages severely and significantly reduced the number of proliferating *pax3a*^+^ muscle stem cells within the injury site, reducing the number of EdU positive cells to homeostatic levels present within uninjured regions (51% decrease in proliferating *pax3a*^+^ muscle stem cells, n=13, Fig. 2n-o). Collectively, these results reveal a surprisingly direct role for a specific macrophage subset in controlling muscle regeneration, demonstrating that a proportion of wound-attracted macrophages form a transient stem cell niche. Ablation of this niche-specific macrophage subset leads to a severe reduction in the number of myogenic progenitors present within the injury site, and a consequent muscle regeneration deficit (Fig. 2f-h).

To establish the nature of injury-responsive macrophage populations, we carried out single-cell RNA-sequencing (scRNA-seq) on injury-located macrophages isolated from discrete phases of the regenerative process. Following needle stick skeletal muscle injury of Tg(*mpeg1*:mCherry) larvae, the wound region was dissected out at 1, 2 and 3 dpi and, mCherry expressing macrophages were FACS sorted (Fig. 3a). Macrophages sorted from uninjured larvae were also included in the analyses. Unsupervised clustering identified seven discrete clusters of macrophage subtypes (Fig. 3b). Of note, cells in all clusters expressed the pan-leukocyte marker *L-plastin* (lymphocyte cytosolic protein 1, *Icp1*) and the pan macrophage marker *cd163*, validating their macrophage identity (Fig. 3e). This analysis revealed greater macrophage heterogeneity than previously described during muscle, or any other regenerative scenario^8-13^ (Fig. 3b). We next compared the time-point at which the injury-responsive macrophages were isolated, versus the identified clusters overlaid on a tSNE plot, to explore if injury times corresponded to specific individual clusters (Fig. 3c). Macrophages from the uninjured larvae formed a temporally uniform cluster (Cluster 3, Fig. 3d). Furthermore, unsupervised clustering of macrophages only from uninjured larvae revealed no further complexity, re-affirming their homogenous nature. This lack of predetermination in the uninjured macrophage population is intriguing given that only a proportion of muscle resident macrophages respond by migrating to the injury. It suggests that the ability to respond to wound derived cues is an acquired state, at least at the level of gene expression. The majority of transient macrophages (1 dpi) also clustered together (Cluster 2, Fig. 3d), suggesting a systematic initial activation of the macrophages that migrate into the injury. The remaining 5 clusters were composed of dwelling macrophages (2-3 dpi), highlighting their heterogeneous nature (Cluster 1, 4, 5, 6 and 7, Fig. 3d). In mammals, wound-associated macrophages have classically been defined as a simple dichotomy between pro-inflammatory, or M1, macrophages and an alternatively activated, anti-inflammatory, or M2^14^, macrophage pool. Although recent *in vivo* analyses clearly illustrate that this classification represents an oversimplification^14^, it provided a starting point to identify and define novel macrophage subtypes *in vivo*. Cells in Cluster 4 were enriched for mammalian M2 markers including arginase 2 (*arg2*), matrix metalloproteinase 9 (*mmp9*) and matrix metalloproteinase 13 (*mmp13*))^15-17^, suggesting a dwelling macrophage-specific anti-inflammatory subtype (53.5%, 100% and 98.6% of Cluster 4 present cells express *arg2, mmp9* and *mmp13*, respectively, Fig. 3e, Extended Data Fig. 6). Since all Cluster 4 cells expressed *mmp9*, we assessed its spatiotemporal expression following injury and observed spatial restriction to the wound site and up-regulation from 2 dpi onwards in the muscle stem cell-associated dwelling macrophage population (Fig. 3f-h). These analyses indicate that Cluster 4 contains a specific muscle stem cell-associated dwelling macrophage subset.

We next examined the list of genes encoding secreted pro-mitogenic molecules specifically up-regulated within Cluster 4 compared to all other identified clusters (Extended Data Fig. 6). This was matched with data sets comprising receptors known to be expressed on muscle stem/progenitor cells^18-20^, leading to the identification of a pairing between nicotinamide phosphoribosyltransferse (NAMPT/Visfatin/PBEF) and the chemokine receptor CCR5 (Fig. 3e). NAMPT is a multi-functional protein, which carries out a well-characterised role in cellular metabolism and NAD regeneration^21^ when localised intracellularly. In addition, its secreted form (secreted NAMPT (secNAMPT)) has been documented to function as a cytokine with reports of both physiological and pathological functions in a number of tissues^22^. In contrast to its intracellular function the role of the secreted protein is less well understood, with respect to both its mode of secretion as well as mechanism of action. Despite this lack of mechanistic insight, secNAMPT has been demonstrated to be pro-mitogenic for a wide variety of cell types, and is also a positive regulator of a broad variety of regenerative capacities in distinct tissues^22^. CCR5 has been suggested as a putative cell surface receptor for NAMPT^23^. Although this premise is supported by only a single report, this study did demonstrate the direct binding of NAMPT to CCR5 by surface plasmon resonance^23^. Furthermore, following mouse muscle injury, there is a temporally restricted up-regulation of CCR5 in the regenerate during the injury resolution phase where muscle stem cell proliferation is documented to occur^19^ and experiments on cultured myoblasts have shown an inhibition of proliferation upon antibody-mediated blocking of CCR5^18^. Additionally, NAMPT has been demonstrated to alter the expression of myogenic regulatory factors in myoblasts *in vitro*^24^.

To examine this possibility, we determined the spatio-temporal expression of *nampt* within wound dwelling macrophages *in vivo* using a number of independent approaches. Firstly, we undertook *in situ* hybridisation analyses post-larval muscle injury and observed that *nampt* was specifically upregulated within the wound site at 2 and 3dpi (Fig. 4a). Nampt antibody staining was consequently performed in the compound Tg(*mpeg1*:GAL4FF/*UAS*:NfsB-mCherry);TgBAC(*pax3a*:GFP) transgenic line, confirming that at 2 dpi Nampt expression was localised specifically to the stem cell-associated dwelling macrophages as well as the extracellular space of the injury site (Fig. 4b). A third approach made use of the key physiological role of intracellular NAMPT in catalysing the rate-limiting process in the NAD salvage pathway^25,26^. Increased NAMPT activity has been documented to elevate levels of intracellular NAD^+^/NADH^27^. NADH exhibits endogenous fluorescence with an excitation peak at about 365 nm that is amenable to examination by two-photon excitation fluorescence microscopy^28,29^. We therefore developed an assay to examine NADH levels *in vivo*, and used this as a proxy to estimate cellular Nampt levels in real time within the wound milieu. At 2 dpi, dwelling macrophages alone specifically expressed high levels of NADH within the wound site (Fig. 4c), an observation that is indicative of increased Nampt production specifically within the dwelling macrophages and highlighting their role as the primary source of Nampt in the wound site.

We next sought to establish if secNAMPT acted to regulate muscle stem cell proliferation through its putative receptor, CCR5. Since there is only a single report of a direct NAMPT-CCR5 interaction, we set out to independently evaluate NAMPT binding to CCR5. The binding of human recombinant NAMPT (hrNAMPT source 1 (hrNAMPT_(1)_)) to recombinant CCR5 receptor (hrCCR5) was evaluated via an enzyme-linked immunosorbent assay (ELISA) approach. We found that hrNAMPT_(1)_ binds hrCCR5 with a dissociation constant (K_D_) of 172±18 nM (Extended Data Fig. 7), confirming previous findings^23^. Next, we utilised *in vitro* cell culture models to determine if NAMPT initiates a pro-proliferative signalling cascade on muscle stem cells and assess if the NAMPT-CCR5 interaction yields functional modulation of this process. Initially, two human recombinant NAMPT protein sources (hrNAMPT_(1)_ and hrNAMPT_(2)_) were applied to mouse C2C12 myoblasts and proliferation assayed by means of EdU incorporation. Both sources of NAMPT resulted in comparable and significant dose dependent increases in myoblast proliferation (Extended Data Fig. 8a). This proliferative response was blocked in the presence of the dual CCR2/CCR5 antagonist, cenicriviroc (CVC)^30^ and CCR5 selective antagonist maraviroc (MVC)^31^, but not in the presence of the CCR2 selective antagonist PF-4136309 (PF)^32^ (Extended Data Fig. 8a), highlighting selective signalling of NAMPT via CCR5 to induce myoblast proliferation.

Next, we turned to an *in vitro* multi-cell co-culture system to examine the relationship between macrophages and NAMPT-mediated proliferation. We assessed the proliferation of isolated embryonic mouse PAX7^+^ muscle stem cells in three culture conditions: embryonic mouse primary myoblast only (MB), and co-cultures of mouse myoblast with either mouse macrophages (MB+MΦ) or 3T3 mouse fibroblast cells (MB+3T3). The mouse macrophage cell line utilised is known to express high levels of NAMPT^33^, while 3T3 fibroblast cells do not naturally secrete NAMPT in culture^34^. In these experiments, we observed that the presence of macrophages promoted satellite cell proliferation (Fig. 4d), a response not elicited when macrophages were replaced in the co-culture by 3T3 cells (Fig. 4d). Furthermore, addition of hrNAMPT was able to increase the proliferation of satellite cells in MB and MB+3T3 culture conditions to a comparable level to that evident in MB+MΦ cultures (Fig. 4d), an effect that was abolished in the presence of CVC (Fig. 4d). Addition of CVC also reduced satellite cell proliferation in MB+MΦ culture conditions to levels evident in MB alone or MB+3T3 cultures (Fig. 4d), highlighting that the macrophage driven increase in satellite cell proliferation was primarily mediated by secNAMPT. Interestingly, hrNAMPT did not increase satellite cell proliferation in the presence of macrophages suggesting that there is a maximum threshold rate of proliferation and this may be modulated by receptor saturation. Collectively, these observations support the hypothesis that macrophage secreted NAMPT is a critical pro-proliferative signal that activates the CCR5 receptor present on satellite cells to stimulate muscle regeneration.

Subsequently, we sought to validate these findings *in vivo*. We confirmed that the zebrafish *ccr5* orthologue was expressed on 2 dpi FACS sorted *pax3a*^+^ myogenic stem/progenitor cells by RT-PCR (Extended Data Fig. 9a-b), and administered CVC and MVC to larval zebrafish to block Ccr5 signal propagation following needle stick muscle injury. Using birefringence imaging we detected a highly significant regeneration deficit in the drug-administered larvae (Fig. 4e-f). This was not due to effects on macrophage migration kinetics or a block in macrophages transitioning from a transient to a dwelling state, both of which are phenotypically wild-type upon drug administration (Extended Data Fig. 9c-f). Rather, we observed a reduction in the proliferation of *pax3a*^+^ myogenic stem cells present in the wound site that was similar in magnitude to that observed post-ablation of dwelling macrophages (Fig. 4g-h). Collectively, these data are consistent with Ccr5-dependent signalling being required for muscle stem cell proliferation during regeneration *in vivo*.

Unconditional NAMPT gene knockout in mice results in embryonic lethality^35^. To circumvent this eventuality confounding our analyses, we developed a macrophage specific loss-of-function mutagenesis system. We generated stable macrophage-specific expression of Cas9 from a Tg(*UAS*:NLS-Cas9) transgene coupled to a Tg(*mpeg1*:Gal4FF) transgene. By delivering a *nampt* guide RNA (gRNA) by microinjection, we achieved durable macrophage specific *nampt* gene editing and subsequently assayed macrophage-derived *nampt’s* role in muscle repair (Fig. 4i). Immunostaining for Nampt revealed a visible reduction in Nampt expressing cells present in the wound site following needle stick muscle injury in the *nampt* gRNA injected larvae (Extended Data Fig. 10a). Furthermore, quantification of NAD^+^/NADH levels in isolated macrophages functionally validated macrophage specific *nampt* loss-of-function using this macrophage specific gene editing approach (Extended Data Fig. 10b).

Following needle stick muscle injury, *nampt* gRNA injected larval macrophages responded by migrating to the injury zone and locating within the wound boundaries from 1 to 3 dpi (Extended Data Fig. 10c). However, these *nampt*-deficient macrophages failed to induce appropriate cell proliferation and regeneration at the injury site (Fig. 4j-m), highlighting a specific requirement for functional macrophage-derived Nampt to ensure appropriate regeneration. Collectively, our *in vitro* cell culture analyses, together with our *in vivo* chemical inhibition and macrophage-specific *nampt* loss-of-function studies demonstrate, a requirement for macrophage-derived NAMPT to stimulate muscle stem cell proliferation in a CCR5-dependent manner (Fig. 4n).

The satellite cell is archetypal of a unipotent tissue resident stem cell that occupies a specific anatomical niche within a differentiated tissue. Decades of research have revealed the extraordinary capacity of this system to effectively coordinate muscle repair in response to a wide variety of insults. Despite this demonstrated regenerative capacity, transplantation of isolated muscle stem cells has yet to provide therapeutic impact, and pro-regenerative treatments that stimulate muscle stem cells are entirely lacking at this juncture. Our data suggest that even this simplest of stem cell systems requires a complex interaction with a range of cellular contexts during repair *in vivo*, and that the innate immune system, in particular the macrophage, is a key modulator of the regenerative process. We introduce the concept of a specific macrophage subset, one amongst a surprising complexity of macrophage phenotypes present during wound repair, acting as a transient stem cell niche that directly regulates muscle stem cells through the provision of mitogenic stimuli, specifically the NAMPT/CCR5 axis. Collectively, our results suggest that providing specific macrophage-derived signals required for muscle stem cell proliferation, such as the ones we identified here, may provide an avenue to achieve better myoblast-based therapy outcomes.

## Methods

### Zebrafish strains and maintenance

Existing transgenic lines used were, TgBAC(*pax3a*:GFP)^i150^ (referred to as TgBAC(*pax3a*:GFP))^36^, Tg(*mpeg1*:mCherry)^gl23^ (referred to as Tg(*mpeg1*:mCherry))^37^, Tg(*mpeg1*:GAL4FF)^gl25^ (referred to as Tg(*mpeg1*:GAL4FF))^37^, Tg(*UAS-E1b*:Kaede)^s1999t^ (referred to as Tg(*UAS*:Kaede))^38^, Tg(*UAS-E1b*:Eco.NfsB-mCherry)^c264^ (referred to as Tg(*UAS*:NfsB-mcherry)^39^, Tg(−8*mpx*:KALTA4)^gl28^ (referred to as Tg(*mpx*:KALTA4))^40,41^, Tg(*actc1b*:EBFP2)^pc5^ (referred to as Tg(*actc1*:BFP))^42^, Tg(*ubi:*secAnnexinV-mVenus)^mq8Tg^ (referred to as Tg(*ubi*:secAnnexinV-mVenus))^43^ and Tg(*actc1b*:GFP)^zf10^ (referred to as Tg(*actc1b*:GFP))^44^. All experiments were conducted in accordance with Monash University guidelines and approved by the local ethics committee. All procedures involving animals at the Hubrecht Institute were approved by the local animal experiments committees and performed in compliance with animal welfare laws, guidelines and policies, according to national and European law. Staging and husbandry were performed as previously described^45^. All embryos were maintained in Ringer’s solution at 28.5°C and treated with 0.003% 1-phenyl-2-thiourea (PTU) (Sigma-Aldrich) from 8 hpf.

### Muscle injury

4dpf larvae were anaesthetized in 0.01% tricaine (MS-222) (Sigma-Aldrich) in Ringer’s solution. Mechanical injures were targeted either to the dorsal or ventral myotome above the cloaca when the larvae is oriented dorsal to the top, anterior to the left. Needle-stick injury was carried out as previously described^3^. Briefly, the myotome was subjected to a single 30-gauge needle puncture that generates an extensive injury with many damaged muscle fibres. For laser-induced injury, anaesthetised larvae were mounted in a thin layer of 1% low-melt agarose in Ringer’s solution. Injuries were carried out using a UV-nitrogen laser pulsed through a coumarin 440 nm dye cell coupled to a Zeiss Axioplan microscope (MicroPoint Laser System, Andor Technology). On average a laser injury required pulses for 5-10 sec from laser beams focused through a X40 water immersion objective. For time-lapse analysis of the immediate response to injury, muscle fibre ablations were achieved using a SIM scanner (Olympus) at 790 nm and 200 msec dwell time at 100% laser power on an Olympus FVMPE-RS upright multi-photon microscope equipped with a 25X/1.05 water immersion objective. Injury-responding macrophages were tracked using the manual tracking plugin in Fiji^46^.

### Microscopy and image analysis

Whole larva time-lapse imaging to track injury-responding macrophages were performed using a Zeiss Lightsheet Z.1 microscope with a 5X/0.16 air objective and environmental controls (28.5°C). The XY resolution was 1.14 μm, the Z resolution was 5.5 μm, with a lightsheet thickness of 11.68 μm. Total imaging time per larvae was 25 h (1000 3D stacks acquired at 1.5 min intervals), and viability of larvae was confirmed at the end of the imaging session by assessing heart rate. For tracking, macrophage images were first filtered with 3D median filter and then segmented with hysteresis thresholding using algorithms from the 3D ImageJ Suite^47^. Low and high threshold for hysteresis were chosen visually. The datasets were then time reversed, in order to track cells exiting the injury site. The margins of the wound were manually labelled within ImageJ. The tracking procedure was based on overlapping segmentation of cells between consecutive frames. Cases where two cells were too close and formed one object were designated “merging” as the segmentation algorithm was unable to separate them. Tracked macrophage images were reversed for the visualisation of results.

Line-scanning confocal microscopy for long-term time-lapse imaging and single Z-stack acquisition was performed using a Zeiss LSM 710 upright confocal equipped with a 20X/1.0 water immersion objective. Photoconversions were carried out using the bleaching tool with a 405 nm diode laser. Time-lapse imaging at high temporal and spatial resolution was performed on an inverted LSM 880 fast AiryScan confocal equipped with a 40X/1.3 oil immersion objective and piezo Z-stage. The voxel size was kept constant at 0.2 x 0.2 x 1 μm and depending on the field of view frame rates of 3-18 frames per second were achieved. Photobleaching was assayed post imaging and determined to be minimal for imaging durations of up to 1h.

Fixed and Immunostained cell culture samples were imaged on a Leica DMi8 inverted widefield microscope with a X10 objective. Birefringence imaging was carried out as previously described^4^, using a Leica DM IRB upright microscope integrated with the Abrio software (CRI Hinds Instruments) using a X5 objective. Images were analysed using the software Fiji^46^, whereby the mean grey value of the injury site birefringence is normalised to the region before and after it to calculate relative birefringence of the wound site.

Microscopy images were processed in Adobe Creative Cloud 2018, Fiji^46^ and Imaris 9.2 (Bitplane). Counting macrophage numbers and further 3D analyses were performed by surface rendering their volumes using Imaris. Sphericity analysis assessed a cells shape deviation from a perfect sphere, which is assigned an arbitrary value of one. Sphericity values were generated as a summary statistic of the surface render. Proliferating stem cell counts were carried out on Fiji. The PAX7 and EdU acquisition channels were segmented using the threshold command. The image calculator function was used to generate a masked channel of only EDU positive PAX7 cells. The number of cells was counted using the Analyse Particles command.

### Chemical treatments

Cell ablation was carried out as previously described^48^ with minor modifications. Larvae at the appropriate stage were incubated in 5-10 mM metronidazole (Mtz) (Sigma-Aldrich) in Ringer’s solution and daily refreshed until experimental end point. Drug treatments were carried out by incubating 4 dpf needle stick injured larvae in 5 and 10 μM cenicriviroc (CVC) (Med Chem Express) and 5 and 10 μM maraviroc (MVC) (Med Chem Express) in Ringer’s solution immediately post injury and refreshed daily. Cell culture drug supplementation was carried out by adding the following to the growth media of C2C12 cells for 6 h: 1.9, 9.5 and 19 nM recombinant human visfatin (hrNAMPT_(1)_) (PeproTech), 1.9 and 9.5 nM recombinant human visfatin (hrNAMPT_(2)_) (Enzo Life Sciences), 100 nM CVC (Med Chem Express), 100 nM MVC (Med Chem Express) and 100 nM PF-4136309 (PF) (Med Chem Express). The following were added to the proliferation media of primary mouse myoblast co-cultures for 24 h, 9.5nM hrNAMPT_(2)_ (Enzo Life Sciences), and 100 nM CVC (Med Chem Express). LysoTracker assay: Larvae were incubated in 10 μM LysoTracker™ Deep Red (Thermo Fisher) in Ringer’s solution for 1 h in the dark and rinsed 5 times with fresh Ringer’s before imaging.

### 5’-ethnyl-2’-deoxyuridine (EdU) labelling

Larvae: Labelling and detection was carried out as previously described^49^ with minor alteration. 6 dpf/2 dpi larvae were pulsed with 2.5 nM EdU (Thermo Fisher) for 1h and chased for a further 1.5 h prior to fixation. Cell culture: Cells were incubated for 1 h in media supplemented with 10 μM EdU. Following Edu pulse, C2C12 cells were fixed immediately, while primary mouse myoblast co-cultures were rinsed in PBS, replaced with media and incubated for a further 2 h following which the cells were fixed. Cells were processed using the Click-iT™ EdU Alexa Fluoro™ 647 imaging Kit (Thermo Fisher) following the manufacturer’s protocol.

### Immunohistochemistry and *In situ* hybridisation

Antibody staining on whole-mount larvae was carried out as previously described^50^ and on cultured myoblasts as previously described^51^. Following *in situ* hybridisation, antibody staining was carried out using standard procedures. Antibodies: mouse anti-Pax7 (1:10, DSHB), chicken anti-GFP antibody (1:500, Thermo Fisher), mouse anti-mCherry antibody (1:500, Abcam), rat anti-mCherry antibody (1:500, Kerafast) (immunohistochemistry), rabbit anti-PBEF1 (anti-NAMPT) antibody (1:50, Sigma-Aldrich) and secondary Alexa Fluro-coupled antibodies (Thermo Fisher). Nuclei were visualised by staining with DAPI (Sigma-Aldrich). *In situ* hybridisation and probe generation was performed as previously described^52^. Antisense probe used: *mmp9*^53^. *nampt* (ENSDARG00000030598) PCR probe containing a T7 RNA polymerase promoter at the 3’ for the antisense probe and an SP6 RNA polymerase promoter at the 5’ for the sense probe was generated using primers 5’-GAGtatttaggtgacactatagGGTTTCATCGCAAGAGACGG-3’ and 5’-GAGtaatacgactcactatagggGCGGAAGCACCTTATAGCCT-3’.

### NAMPT binding to CCR5

ELISA plates (Medium binding, Greiner Bio-One) were coated with 1% BSA or 20 nM of GST-fused recombinant human CCR5 (hrCCR5, Abcam) in PBS overnight at 4°C. Wells were blocked for 1 h at room temperature with 1% BSA in PBS containing 0.05% Tween-20 (PBS-T). Wells were washed 3 times with PBS-T and further incubated with hrNAMPT_(1)_ (Peprotech) at increasing concentration (0 nM to 800 nM) for 1 h in PBS-T with 0.1% BSA. Bound NAMPT molecules were detected using a biotinylated antibody for NAMPT and HRP-streptavidin (Human PBEF/Visfatin DuoSet ELISA, R&D Systems). Signals obtained on BSA-coated wells were used to remove non-specific binding for each NAMPT concentrations to obtain specific binding values. Specific binding data were fitted by non-linear regression with Prism 7 to obtain the dissociation constant (K_D_) using A_450_ nm = Bmax*[NAMPT]/(K_D_ + [NAMPT]).

### Cell culture

The mouse muscle cell line C2C12^54^ were cultured in growth media (Dulbecco’s Modified Eagle Medium (4.5 g/l D-Glucose, No L Glutamine, No Sodium Pyruvate (Gibco))+20% Fetal Bovine Solution-One Shot (Gibco)+1% Glut Max 100x (Gibco)). Cells were maintained at 37°C, 5% CO_2_. Cells at 70% confluence, passage 8 were extracted from T75 flasks with 0.025% Tryspin EDTA (Gibco), neutralised in growth media, spun at 180 X g for 5 min to pellet cells. The cells were then resuspended in 10 ml of fresh growth media. 500 μl of cells were plated on a 8-well on cover glass II (Sarstedt) chamber slide at a density of 1×10^3^ cells/ml. Cells were left 4 h at 37°C to re-attach. For drug treatments, the media were supplemented with appropriate doses and cultured for 6 h.

For isolation of primary mouse myoblasts, limb skeletal muscle from E17.5 C57/BL6J mice were minced and digested in 0.125% Trypsin at 37°C for 20 min. Fibroblasts were depleted by plating cells in 10 cm^2^ tissue culture dishes (2 embryos per dish) in proliferation media (DMEM + 20%FBS) for 1 h. Media with non-attached cells was re-plated in gelatin-coated 10 cm^2^ tissue culture dishes in proliferation media for 24 h. Myoblasts were again depleted for fibroblasts prior to co-culturing on gelatin-coated 48 well plates in DMEM+20%FBS+10%L929-conditioned medium. 100,000 myoblasts were plated with either 7,500 MafB/c-Maf deficient (Maf-DKO) macrophages^55^ or 1,000 3T3 cells per well. For drug treatments, the media were supplemented with appropriate doses and cultured for 24 h.

### Fluorescence activated cell sorting (FACS), single-cell RNA-sequencing and analysis

Injury-responding macrophages were isolated by FACS as previously described^56^, with the following modifications. 4dpf Tg(*mpeg1*:mCherry) larvae were subject to needle-stick injury and the injured region dissected at 1, 2 and 3 dpi and tissue dissociated into a single cell suspension. Uninjured larval trunk tissue was also included in the analysis. Cells were sorted using a FACS Aria II (BD biosciences). Live individual macrophages (based on mCherry fluorescence, DAPI exclusion and forward and side scatter properties) were sorted into pre-prepared 384 well plates containing 100-200 nl of CEL-seq primers, dNTPs and synthetic mRNA Spike-Ins contained in 5 μl of Vapor-Lock (Qiagen). Immediately following sorting, plates were spun down and frozen at −80°C until sequencing.

Single-cell RNA-sequencing libraries were prepared using the SORT-seq platform as previously described^57^. In this platform the Cel-Seq2 protocol^58^ is followed with the aid of robotic liquid handlers. This protocol results in each cell being barcoded, and generating single-cell transcriptomes of all isolated macrophages. Each time point is independently replicated and results in approximately 768 macrophages per time point (3072 macrophages in total) to be individually sequence. Next generation sequencing was carried out using an Illumina NextSeq platform. Paired reads were mapped against the zebrafish reference assembly version 10 (GRC10). FASTQ files were processed as previously described^57,59^. Paired end read were aligned to the zebrafish transcriptome using bwa^60^ with a transcriptome dataset with improved 3’ UTR annotations to increase the mapability of transcripts^61^. Read 1 was used for assigning reads to correct cells and libraries while read 2 was mapped to gene models. Read counts were first corrected for UMI barcode by removing duplicate reads that had an identical combination of library, cellular and molecular barcodes that were mapped to the same gene. Transcript counts were then adjusted to the expected number of molecules based on counts, 256 possible UMI’s and poissonian counting statistics. The scripts to generate the count files can be found here https://github.com/vertesy/TheCorvinas/tree/master/Python/MapAndGo2 with the readme files found here https://github.com/vertesy/TheCorvinas/blob/master/Python/MapAndGo/Readme_MapAndGo.md.

Spike-in RNAs were discarded and not included in further analysis. Transcript counts of all assayed plates (technical replicates of injury types) were combined into one matrix for downstream analysis. Downstream analysis of the combined samples transcript read counts was performed using Seurat (v2.3.4)^62^. Transcript counts matrix was imported using CreateSeuratObject function (min.cells = 25, min.genes = 250) and low quality cells were discarded with FilterCells with the following thresholds: a minimum of 500 and maximum of 3500 genes, a maximum of 10% of mitochondrial genes and a minimum of 1000 and maximum of 15000 UMIs). The filtered dataset was log-normalised using a scale factor 10000 and a set of ∼2800 genes were used for linear dimension reduction (FindVariableGenes, x.low.cutoff = 0.05, x.high.cutoff = 3, y.cutoff = 0.5). Unwanted sources of variation were removed by regressing out UMI number per cell, percentage of mitochondrial reads per cell. Clustering analysis was performed with Seurat’s FindClusters function (reduction.type = “pca”, dims. Use = 1:50, resolution = 0.5, save.SNN = TRUE) and a non-linear dimensional reduction using RunTSNE (reduction.use = “pca”, dims.use = 1:50, perplexity = 30, tsne.method = “Rtsne”). Seven cell clusters were identified. To identify cluster biomarkers either FindAllMarkers (logfc.threshold = 0.25, test.use = “wilcox”) or FindMarkers (min.pct = 0.20) functions were used to compare all clusters against each other or cluster specific comparisons. tSNE, feature plots and heatmaps were created using Seurat’s TSNEPlot, FeaturePlot and DoHeatmap functions.

### Electron Microscopy

4 dpf larvae were subject to laser ablation skeletal muscle injury. At 22 hpi a confocal stack of the injury region was acquired on a Zeiss LSM 710 upright confocal microscope. Following imaging larvae were euthanized and fixed according standard procedures in 2.5% glutaraldehyde, 2% paraformaldehyde in 0.1 M sodium cacodylate buffer overnight at 4°C. Larvae were next post-fixed with 1% OsO4, 1.5% K_3_[Fe(III)(CN)_6_] and dehydration performed with ethanol and propylene oxide. Larvae were embedded in Epon 812 and ultrathin sections of 70 nm were cut using a diamond knife (Ultra 45° Diatome) on a Leica Ultracut UCT7 and placed on 50 mesh copper grids with carbon coated formvar support film and stained with uranyl acetate and lead citrate. Large area STEM tile sets were taken on a Thermo Fisher/FEI NovaNano SEM 450 equipped with a STEM II (HAADF) detector set at 30 keV, working distance 6.8 mm. MAPS 2.1 software was used to create the tile sets and perform correlations to the confocal data sets. High resolution EM imaging was performed on a Jeol1400Flash TEM at 80 KeV.

### Macrophage specific Nampt loss-of-function mutation

The Tol2-flanked transgene Tg(4x*UAS*:NLS-Cas9, *cryaa*:EGFP)^gl36^ (referred to as Tg(*UAS:*NLS-Cas9)) was assembled by Gateway cloning and microinjected with Tol2 mRNA into the Tg(*mpeg1*:GAL4FF/*UAS*:NfsB-mCherry) background. Co-segregation of the three transgenes through F0 and F1 backcrosses onto this background was achieved by selecting embryos with red macrophages (confirming the presence of the *mpeg1*:GAL4FF transgene) and green eyes (confirming the presence of the *cryaa*:EGFP marker gene linked to the 4x*UAS*:NLS-Cas9). For the macrophage-specific gene editing experiments, embryos from incrossed F2 generation adults were used. The Tg(*mpeg1*:GAL4FF/4x*UAS*:NLS-Cas9,*cryaa*:EGFP*/UAS*:NfsB-mCherry) line is referred to as *mpeg1*-Cas9. Synthetic guide RNAs (gRNAs) targeting *nampt* were generated as crRNA:tracrRNA duplexes (Alt-R® CRISPR-Cas9 system, IDT). Gene-specific crRNA sequences were selected using the Alt-R® CRISPR-Cas9 custom guide RNA design tool (IDT). *nampt*-specific crRNAs were heteroduplexed to universal tracrRNA according to manufacturer recommendations to generate bipartite gRNAs. The *nampt* gRNA efficiencies were intially tested in whole embryo gene editing. Individual guides and recombinant Cas9 protein (Alt-R® S.p Cas9 Nuclease, IDT) were injected into the blastomere of one-cell stage embryos. At 2 dpf, genomic DNA from microinjected embryos was extracted and assayed for CRISPR/Cas9 introduction of genomic editing by PCR using primers flanking the target site. A guide with 100% mutation efficiency (15 individual larvae assayed) was selected for use in tissue specific gene editing (*nampt* crRNA of interest: ACGACAAGACGGTCTTCTAT (GGG)). To induce macrophage specific gene editing 3 nL of gRNA mix (3 μL of 3 μM gRNA + 0.5 μL of 2% phenol red + 1.5 μL 0.1M KCl mix) was injected into the cell of one-cell stage *mpeg1*-Cas9 embryos.

### Visualisation and quantification of NAD/NADH

*In vivo* two-photon excited fluorescence of larval zebrafish NADH was measured on an Olympus FVMPE-RS upright multi-photon microscope using a 25X/1.05 water immersion objective. A wavelength of 810 nm and a 450/70 bandpass filter was used. The same wavelength and a 610/70 bandpass filter were used to detect mCherry fluorescence. A galvo scanner was used to generate high-resolution data sets while an 8 kHz resonant scanner was used for time-lapse imaging to minimise phototoxic effects.

*In vitro* NAD/NADH levels were measured using the NAD/NADH-Glo™ assay (Promega) following supplier instructions. The assay was carried out on macrophage sorted from macrophage specific *nampt* knockout larvae (*mpeg1*-Cas9+*nampt* guide injected) and control (*mpeg1*-Cas9) larvae. At 2 dpf control larvae were soaked in either the NAMPT enzymatic inhibitor GMX1778 (10 μM) (Sigma-Aldrich) or NAMPT’s rate limiting enzyme catalyses product, nicotinamide mononucleotide (NMN) (100 μM) (Sigma-Aldrich). These controls were used as additional controls to identify the assay’s detection range. At 3 dpf *mpeg1*^+^ macrophages from each group were sorted into white, flat bottom 96 well plates (Costar) containing 50 μL PBS at 2000 cells/well density. Cells were incubated in 50 μL of NAD/NADH Glo™ detection reagent. After 1 h incubation, luminescence was determined in a microplate reader (BMG PHERAstar; gain 3600, 1 s integration time). Each point of measurement represents the average luminescence reaction measured in relative luminescence units (RLU).

### RT-PCR

Total RNA from FACS sorted *pax3a*^+^ cells were extracted using TRIzol™ Reagent (Thermo Fisher). cDNA was synthesized using the iScript™ Advanced cDNA Synthesis Kit (Bio-Rad) following manufacturer’s instructions. For RT-PCR, 10 μl reactions were set up using GoTaq® Green Master Mix (Promega). Primers used to amplify zf*ccr5*, 5’-TTATAACCAAGAGACATGTCGGCG-3’ and 5’-ACCCAGACGACCAGACCATT-3’. Primer pair designed to cross a 388 bp intron so that cDNA and genomic DNA (gDNA) templates would result in bands of 191 bp and 579 bp, respectively. The cycling protocol was performed as follows: Initial denaturation at 95°C for 2 min followed by 25 cycles of denaturation at 95°C for 30 sec, annealing at 58°C for 1 min and extension at 72°C for 45 sec followed by a final extension of 5 min at 72°C. Cycling protocol provided allowance to amplify both cDNA and gDNA templates.

### Maximum-likelihood phylogenetic tree analysis

A putative Ccr5 orthologue in zebrafish was identified by BLAST and orthology assessed by phylogenetic analysis. Amino acid sequences were aligned by MUSCLE^63,64^ and trimmed using GBLOCKS^65,66^. PHYML^67^ was used to generate a maximum likelihood tree using the JTT model for protein evolution (as inferred using ProTest v3.4.2)^68^. Trees were visualised using iTOL (v4.3)^69^.

### Statistical analysis

Embryos for each experimental treatment group were assigned randomly and the groups were blinded to the experimenter prior to analysis. All experiments were performed with a minimum of 3 independent biological replicates; exact numbers have been indicated in figures. Statistical analysis was carried out using Prism version 7.0c (GraphPad Software, Inc.). Data was analysed using Student’s unpaired two-tailed t-test when comparing two conditions and Analysis of Variance (ANOVA) with Tukey’s post-hoc analysis when comparing multiple conditions.

## Acknowledgements

We thank the WEHI Centre for Dynamic Imaging, Monash Micro Imaging and Monash Ramaciotti Centre for Cryo-Electron Microscopy for microscopy support, Hubrecht Institute Single Cell Sequencing Facility for sequencing service, Monash FlowCore for sorting of cells using FACS and Monash Fishcore staff for technical assistance. We thank Frank. J. Tulenko for advice on phylogenetic tree construction, Alexandra-Madelaine Tichy and Elliot Gerrard for technical assistance, Benjamin. T. Kile and Pierre Gönczy for comments on the manuscript. P.D.N is supported by an EMBO Long Term Fellowship ALTF1129-2015, HFSPO Fellowship (LT001404/2017-L) and a NWO-ZonMW Veni grant (016.186.017-3). This work was supported by the National Health and Medical Research Council of Australia [P.D.C; APP1041885, APP1136567, APP1104190, APP1159278 to P.D.C. and G.J.L; APP1044754, APP1069284, APP1086020 to G.J.L]. The Australian Regenerative Medicine Institute is supported by grants from the State Government of Victoria and the Australian Government.

## Author contributions

D.R. (live-imaging, photoconversions, immunohistochemistry, *in situ* hybridisation, scRNA-seq sample preparation, macrophage *nampt* loss-of-function studies, RT-PCR, NAD^+^/NADH assay) designed and performed experiments, analysed data and co-wrote the manuscript; P.D.N. (scRNA-seq sample preparation), V.W. (live-imaging), L.A.G. (primary myoblast culture), A.J.W. (C2C12 cell culture), Z.J. (ELISA) and V.O. (CLEM) performed experiments; A.I. (macrophage specific gene mutagenesis system) established technique; F.J.R. (scRNA-seq analysis) and T.B. (Lightsheet cell tracking) analysed data; K.L.R., C.M. and J.B. provided reagents; P.D.C., G.J.L. and M.M.M. designed and supervised experiments and P.D.C. co-wrote the manuscript.

## Author information

Correspondence and requests for materials should be addressed to P.D.C.

## Extended Data Figure legends

**Extended Data Figure 1. Dwelling macrophages establish a transient regenerative niche following needle stick injury. a-c’’’**, Maximum intensity projection images of the same individual fish transgenic for Tg(*actc1*:BFP), labelling differentiated muscle fibres (magenta, **a’-c’**), TgBAC(*pax3a*:GFP), labelling *pax3a*^+^ myogenic cells (cyan, **a’’-c’’**) and Tg(*mpeg1*:GAL4FF/*UAS*:NfsB-mCherry) labelling macrophages (yellow, **a’’’-c’’’**) following needle stick muscle injury. **d**, By 2 dpi 51.6% of original injury-responsive MΦs remain in the wound site, these dwelling MΦs go onto establish a transient injury niche that is pro-myogenic (n=20). Mean±S.D. Significance (\*\*\*\**P*<0.0001) in one-way ANOVA with Tukey’s multiple comparison test.

**Extended Data Figure 2. Establishing nitroreductase-mediated immune cell ablation parameters. a-c**, The transgenic line, Tg(*mpeg1*:GAL4FF/*UAS*:NfsB-mCherry) expresses the enzyme nitroreductase specifically within macrophages, enabling the temporally controlled genetic ablation of macrophages by addition of the pro-drug metronidazole (Mtz). The optimum Mtz concentration was established by treating larvae with 5 and 10 mM Mtz for 24 h and visualising (**a**) and quantifying trunk present MΦs. 77% macrophage ablation was achieved by 5 mM Mtz treatment for 24 h from 4 dpf onwards (**b**, n=10). Although uninjured larvae were able to tolerate 10 mM Mtz, needle stick injury carried out at this concentration resulted in significant lethality. Therefore, all further experiments used 5 mM Mtz for MΦ ablations. (**c**) Titrating the Mtz dosing time to specifically ablate MΦ subsets of interest. The injury-responding MΦs (yellow) are shown superimposed on brightfield images of the zebrafish trunk, DMSO treated larvae (vehicle control) exhibit injury-present MΦs at both 1 (transient MΦs) and 3 dpi (dwelling MΦs). Mtz treatment at the point of injury results in the ablation of all injury-responding MΦs, whereas Mtz treatment at 1.75 dpi (42 hpi) results in the ablation of wound-located dwelling MΦs (n=10). **d-g**, As a control experiment for nitroreductase mediated cell ablation, neutrophils, another major component of the innate immune response, were selectively ablated using the transgenic line Tg(*mpx*:KALTA4/*UAS*:NfsB-mCherry). Ablation efficiency following 24 h of Mtz treatment were visualised (**d**) and quantified (**e**, n=10). (**f**) Following needle stick injury, larvae were soaked in Mtz to ablate neutrophils and regeneration monitored by imaging for birefringence and quantified (**g**, n=12). Neutrophil ablated larvae showed no regeneration defects at either Mtz doses, highlighting the specific requirement of macrophages for regeneration and precluding the possibility that the regeneration defect was metronidazole-related. Furthermore, this finding also suggests that 10 mM Mtz treated larval lethality evident in the macrophage ablated setting cannot be attributed to Mtz induced toxicity, rather it’s more likely due to an inability to tolerate injury when close to 90% of macrophages are ablated. **b**,**e**, Mean±S.D. Significance (\*\*\*\**P*<0.0001, \*\*\**P*<0.001, \*\**P*<0.01) in one-way ANOVA with Tukey’s multiple comparison test. **g**, Mean±S.D. Significance (ns, not significant) in two-way ANOVA with Tukey’s multiple comparison test.

**Extended Data Figure 3. Macrophage ablation does not disrupt tissue debris clearance following muscle injury. a-e**, Following macrophage ablation the wound site is cleared of damaged muscle fibres (**a**, phalloidin staining, n=12), acidic organelles (LysoTracker (magenta) imaging (**b**) and quantification (**c**), n=10) and apoptotic cells (line Tg(*ubi*:secAnnexinV-mVenus) (magenta) imaging (**d**) and quantification (**e**), n=20). **c, e**, Mean±S.D. Significance (\*\*\*\**P*<0.0001, ns, not significant) in unpaired *t*-test (**c**) or in two-way ANOVA with Tukey’s multiple comparison test (**e**).

**Extended Data Figure 4. Dwelling macrophage-muscle stem cell associations precede muscle stem cell proliferation. a-b’**, Muscle stem cells migrate into the injury site independent of macrophage derived signals. As macrophages (Tg(*mpeg1*:GAL4FF/*UAS:*NfsB-mCherry), yellow) and muscle stem cells (TgBAC(*pax3a*:GFP), cyan) migrate and populate the injury site simultaneously, we assessed if there was a dependence of one on the other. Both control larvae (**a**) and macrophage ablated larvae (**b**) displayed *pax3a*^+^ stem cells in the injury site at 2 dpi (arrowheads). In the control setting these cells were associated with macrophages. At 3 dpi, control larvae displayed regenerated cyan fluorescence-persistent muscle fibres that arose from *pax3a*^+^ stem cells present in the wound (**a’**), a hallmark of a healing muscle injury. In contrast, although the wound site was still occupied by *pax3a*^+^ stem cells at 3dpi in the macrophage ablated larvae, there were no nascent *pax3a*^+^ stem cell derived muscle fibres present (**b’**, arrowheads), highlighting that macrophage presence in the wound site is not prerequisite for *pax3a*^+^ myogenic cell migration, however, it is required for those migrated cells to enter the myogenic replenishment program (n=12) **c**, Dwelling macrophages interact with wound-responsive muscle stem cells. Examples of 3 independent laser ablation muscle injury sites showing dwelling macrophage-stem cell associations followed by stem cell divisions (arrowheads). Imaging carried out in transgenic zebrafish larvae, Tg(*mpeg1*:GAL4FF/*UAS:*NfsB-mCherry), labelling macrophages (yellow) and TgBAC(*pax3a*:GFP) labelling *pax3a*^+^ cells (cyan). Frames extracted from Supplementary Video. 7.

**Extended Data Figure 5. Correlative light and electron microscopy (CLEM) analyses of the transient macrophage-stem cell niche reveals that macrophages and stem cells maintain direct hetero-cellular surface appositions in x-y-z planes. a**, Confocal microscopy of the wound site at 25 hours post laser ablation muscle injury in the compound transgenic zebrafish line, Tg(*mpeg1*:GAL4FF/*UAS*:NfsB-mcherry); TgBAC(*pax3a*:GFP), labelling macrophages (yellow) and *pax3a*^+^ muscle stem/progenitors (cyan). **b**, Large STEM tile set of the same trunk region of the identical larvae illustrated in **a** generated after Epon-embedding and sectioning. This data set was used to correlate and identify the highlighted macrophage-stem cell interaction of interest (asterisk (**a**,**b**)). Dotted square marks the area that was further examined by transmission electron microscopy (in **c**,**d**). **c**, Region of interest encompassing a macrophage and stem cell which maintain a close interaction examined through z-depth. The cells of interest and interaction area are segmented (macrophage cytoplasm; yellow, macrophage nucleus; orange, stem cell cytoplasm; cyan, stem cell nucleus; blue, cell-cell interaction surface; magenta). **d**, high resolution images of two planes through the z-depth (plane shown correlated with red dotted lines to the segmentation images) further demonstrate the close association between the two cells.

**Extended Data Figure 6. Differentially expressed genes in macrophage Cluster 4 as identified by scRNA-seq**. Table recording the top 100 differentially expressed genes in macrophages located in tSNE Cluster 4 verses all other sequenced macrophage clusters.

**Extended Data Figure 7. NAMPT binding to CCR5**. ELISA plates were coated with CCR5 or BSA and further incubated with hrNAMPT_(1)_ at increasing concentration (0 nM to 800 nM). NAMPT molecules bound to CCR5 were detected via a biotinylated antibody. NAMPT binds to CCR5 with a K_D_ of 172±18 nM (n=2 in triplicate). The graph shows a representative binding curve where non-specific binding to BSA was deducted. Means±S.E.M.

**Extended Data Figure 8. Exogenous NAMPT supplementation can enhance myoblast proliferation *in* vitro. a**, *In vitro* assay assessing the effects of exogenously introduced factors on C2C12 myoblast proliferation. Proliferation is identified by EDU incorporation. NAMPT administration (2 commercially available NAMPT sources tested, hrNAMPT_(1)_ and hrNAMPT_(2)_) leads to a dose dependent increase in myoblast proliferation. This effect is specifically mediated via the CCR5 receptor. Co-administration of NAMPT with the CCR2/CCR5 dual inhibitor cenicriviroc (CVC) and CCR5 inhibitor maraviroc (MVC) abolishes NAMPT’s pro-proliferative response, while co-administration with the CCR2 inhibitor PF-4136309 (PF) does not hinder NAMPT’s stimulatory effect on myoblast proliferation. Total number of nuclei assayed for each group recorded on the graph. Mean±S.D. Significance (\*\*\*\**P*<0.0001, \*\**P*<0.01) in two-way ANOVA with Tukey’s multiple comparison test.

**Extended Data Figure 9. Inhibiting Ccr5 receptor activation does not affect injury-responsive macrophage dynamics. a**, The predicted orthologous *Ccr5* gene in zebrafish was identified by determining the appropriate best hit for the human CCR5 amino acid sequence in the zebrafish genome using BLAST. A maximum-likelihood phylogenetic tree constructed using protein sequences of Ccr5 positioned the putative zebrafish Ccr5-like (Ccr5L) sequence as a homolog of mammalian CCR5 with strong bootstrap support. Bootstrap values with 500 replicates are documented below branches. CCR7 was used as an out-group. The accession numbers for genes included in our analysis are provided in the table on the right. **b**, *pax3a*^+^ muscle stem cells express zfCcr5 receptor at 2 dpi as detected by RT-PCR. **c-d**, Ccr5 inhibition by the receptor antagonist cenicriviroc (CVC) does not affect macrophage migration into the injury site nor the successful transition into a dwelling MΦ subtype. (**d**) Quantification. **e-f**, CVC treated macrophages appear morphologically indistinguishable from the control with the transient MΦ having lower sphericity values then their dwelling counterpart. (**f**) Quantification. **d**,**f**, Mean±S.D. Significance (\*\*\*\**P*<0.00001, ns, not significant) in two-way ANOVA with Tukey’s multiple comparison test.

**Extended Data Figure 10. Validating the macrophage specific *nampt* gene editing strategy. a**, Nampt protein expression in the wound site was assessed in the *nampt* gRNA injected *mpeg1*-Cas9 larvae following needle stick muscle injury. The gene-edited larvae presented observably reduced Nampt expression (magenta) within the injury zone (n=12). **b**, FACS isolated macrophages from gene-edited larvae were assayed for Nampt functionality by measuring NAD/NADH levels using a luminescence based assay. Macrophages isolated from control-uninjected larvae were used to measure the baseline NAD/NADH levels for larval zebrafish macrophages. Macrophages from larvae treated with the Nampt enzymatic inhibitor GMX1778 was used to identify the NAD/NADH levels present in macrophages in the absences of Nampt function. Furthermore, macrophages from larvae treated with NMN, the main product of Nampt’s rate-limiting enzymatic reaction was used to establish the maximum threshold of the assays sensitivity. Macrophages isolated from *nampt* gRNA injected *mepg1-*Cas9 larvae presented with significantly reduced levels of NAD/NADH, reflective of a functional loss-of-function of Nampt in the macrophages of these larvae. Mean±S.D. Significance (\*\*\*\**P*<0.00001, \*\*\**P*<0.001) in one-way ANOVA with Tukey’s multiple comparison test. **c**. Macrophage dynamics of the *nampt* gene edited larvae were assayed post-wounding and demonstrated no observable deviation from the control-uninjected larvae (n=10).

## Supplementary video legends

**Supplementary video. 1. Following injury macrophages in the vicinity of the wound site mount an immediate wound migratory response**. The video begins by showing a 4 dpf zebrafish larva’s unperturbed skeletal muscle (Tg(*actc1b*:GFP), magenta) with patrolling muscle resident macrophages (Tg(*mpeg1*:GAL4FF/*UAS*:NfsB-mCherry), yellow). 3 frames into the video the larva is subject to an acute laser ablation injury targeting the ventral myotome. Following injury, a subset of macrophages near the wound site actively migrates into the injury site (see stills in Fig. 1a-b’’). Migrating macrophages project large cytoplasmic protrusion on route to the wound site. Macrophages migrate along the mid line as well as over both the dorsal and ventral myotome. Each migrating macrophage path is shown in individual colours (see stills Fig. 1c). Maximum intensity projections of confocal z-stacks imaged at each time point. Larval view: lateral, anterior to left. Video length: 1.02 h, 4 frames/min. Video representative of n=4 observations.

**Supplementary video. 2. A subset of injury-responding macrophages ‘dwell’ in the wound site for the duration of the regenerative process**. Video shows two clips presented side-by-side extracted from a single continuous 24 h confocal time-lapse movie of macrophages (Tg(*mpeg1:*GAL4FF/*UAS*:NfsB-mCherry), yellow) responding to skeletal muscle injury (Tg(*actc1b*:GFP), magenta). The clip on the left shows the early macrophage response to injury from 2 to 9 hours following injury. Macrophages migrate into the wound site extending long cytoplasmic protrusions. There is an initial increase in the wound-located macrophages. These transient macrophages remain in the wound site up to 9.80 h (see stills in Fig. 1d). The clip on the right shows the late injury-present dwelling macrophages, which are composed of 50% of the original macrophages that migrated into the wound site (see stills in Fig. 1e). Dwelling macrophages remain in the wound site until injury resolution is complete (see macrophage injury residing time quantification in Fig. 1f). Maximum intensity projections of confocal z-stacks imaged at each time point. Larval view: lateral, anterior to left, laser ablation targeted to ventral myotome. Video length: left clip 7 h, right clip 6 h, 6 frames/h. Video representative of n=8 observations.

**Supplementary video. 3. Transient and dwelling macrophages are phenotypically distinct**. This video illustrates the morphological differences between early wound-located transient macrophages and late wound remaining dwelling macrophages. The video is presented in a tabular format with the left column highlighting a transient macrophage and the right a dwelling macrophage. In the rows, the bottom row shows the fluorescent pre-processed image of the macrophage (Tg(*mpeg1:*GAL4FF/*UAS*:NfsB-mCherry), yellow) while the top row shows the processed surface rendered image of the macrophage. The surface rendering is pseudocoloured with a sphericity scale where a perfect sphere would have a sphericity value of 1. Transient macrophages have lower sphericity values while dwelling macrophages appear more rounded having significantly higher sphericity values (see graph in Fig. 1i-j).

**Supplementary video. 4. Macrophages that go onto dwell are initially located in close proximity to the wound site at the point of injury**. Lightsheet microscopy time-lapse imaging video of whole 4 dpf Tg(*mpeg1:*GAL4FF/*UAS*:NfsB-mCherry)/Tg(*actc1b*:GFP) (grey) zebrafish larvae documenting macrophage response to laser ablation injury from 0.17 h post laser ablation injury until 19.04 h post injury. Retrospective tracking identified macrophages that go onto dwell within the wound site and the video was processed such that these macrophages are highlighted in yellow and their migration into the injury site (blue) can be observed over the imaged time course. Maximum intensity projection of confocal z-stacks imaged at each time point. Lateral view: anterior to the right, dorsal to the top. Video length: 19 h, 0.67 frames/min. Video representative of n=4 observations.

**Supplementary video. 5. Once macrophages transition to a dwelling state they actively start interacting with *pax3a***^**+**^ **muscle stem cells present in the wound site**. Video consists of two clips extracted from two time-lapse movies taken in immediate succession of the same larvae revealing macrophages (Tg(*mpeg1:*GAL4FF/*UAS*:NfsB-mCherry), yellow) and *pax3a*^+^ muscle stem/progenitor cells (TgBAC(*pax3a*:GFP), cyan) responding to muscle injury (muscle injury shown in differential contrast). The first clip is immediately following injury and reveals that both macrophages and *pax3a*^+^ cells migrate into the injury site independently of each other. *pax3a*^+^ cells from the undamaged regions of the ablated myotome as well as originating from adjacent myotomes migrate to line the edge of the wound site (*pax3a*^+^ cells that go onto line the wound edge are indicated by magenta arrowheads) (see stills in Fig. 2i-k). At the beginning of the second clip (10.25 hpi) macrophages have transitioned to a dwelling state. Dwelling macrophages initiate intense and continuous interactions with *pax3a*^+^ cells lining the wound edge (see stills in Fig. 2l). A surface rendered image of these interactions is shown to further highlight the specific nature of these associations. Maximum intensity projections of confocal z-stacks imaged at each time point. Larval view: lateral, anterior to left, laser ablation to ventral myotome. Video length: 1^st^ clip 4 h, (1.54 frames/min), 2^nd^ clip 3.75 h (0.5 frames/min). Video representative of n=5 observations.

**Supplementary video. 6. Following dwelling macrophage interactions associated myogenic cells undergo cell division**. Video shows an example of an injury-present dwelling macrophage (Tg(*mpeg1:*GAL4FF/*UAS*:NfsB-mCherry), yellow) interacting with a *pax3a*^+^ cell (TgBAC(*pax3a*:GFP), cyan) which consequently undergoes cell division (see stills in Fig. 2m). The video shows in high resolution the spatial control macrophages exert on specific myogenic cells present in the wound site. Maximum intensity projections of confocal z-stacks imaged at each time point. Video commences at 20 hpi. Larval view: lateral, anterior to left, laser ablation to ventral myotome. Video length: 0.65 h, 14.75 frames/min. Video representative of n=10 time-lapse videos documenting n=26 cell divisions.

**Supplementary video. 7. Further documentation of dwelling macrophage associated *pax3a***^**+**^ **cells undergoing cell division**. Video showing dwelling macrophage-myogenic cell contacts leading to the associated myogenic cell undergoing cell division in 3 independent larval laser ablation injuries. Maximum intensity projections of confocal z-stacks imaged at each time point. White arrowheads point to cell undergoing division. Larval view: lateral, anterior to left. Clip 1 (see stills in Extended Data Fig. 5a, injury 1), video commences at 18.5 hpi, video length: 0.4 h, 15 frames/min. Clip 2 (see stills in Extended Data Fig. 5a, injury 2), video commences at 17 hpi, video length: 0.32 h, 7.25 frames/min. Clip 3 (see stills in Extended Data Fig. 5a, Injury 3), video commences at 26 hpi, video length: 5.91 h, 1.4 frames/min. Video’s representative of n=10 time-lapse videos documenting n=26 cell divisions.

