## Supplementary material for "Macrophages provide a transient muscle stem cell niche via NAMPT secretion": 1

### Extended Data Figure 1

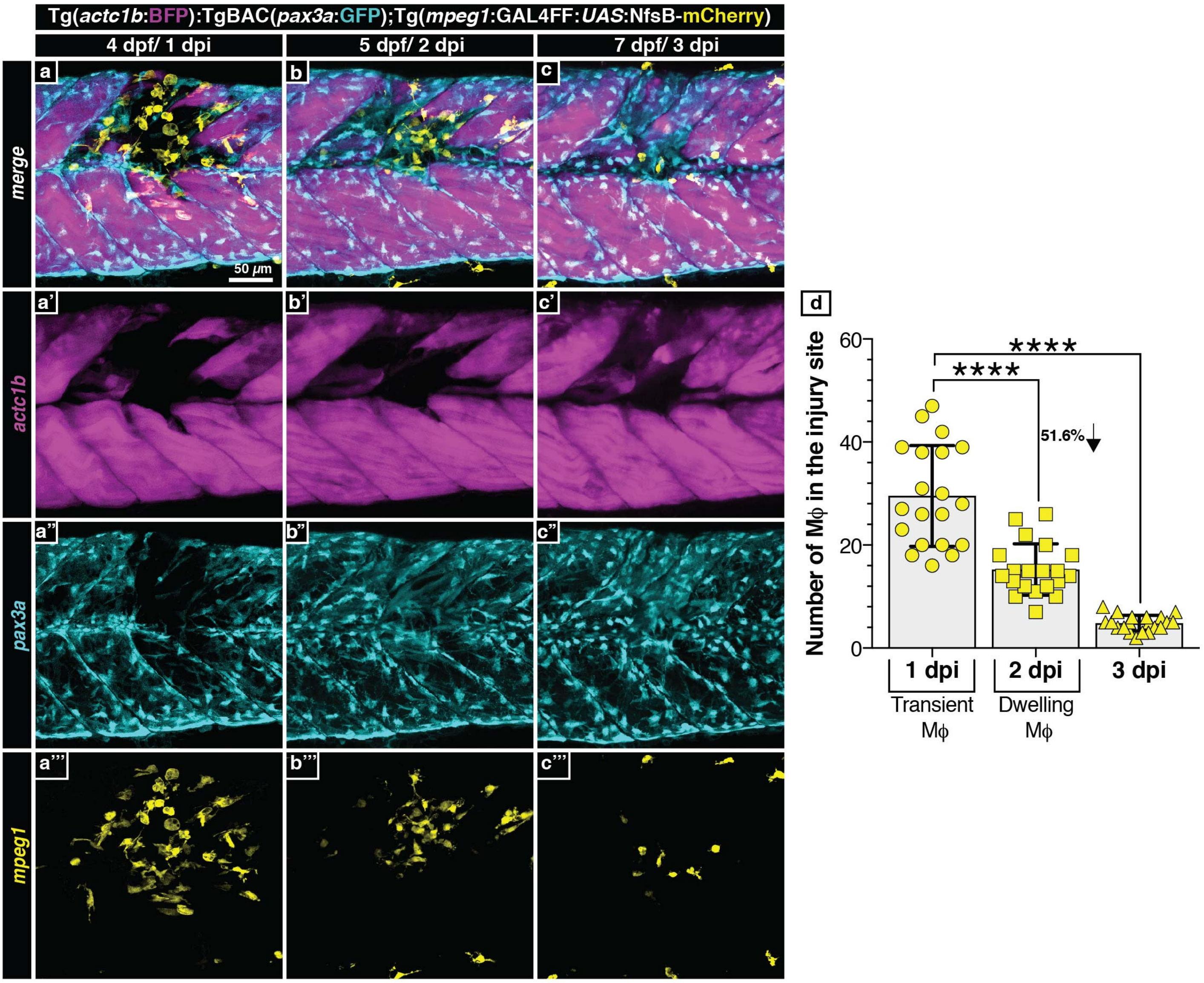

### Extended Data Figure 2

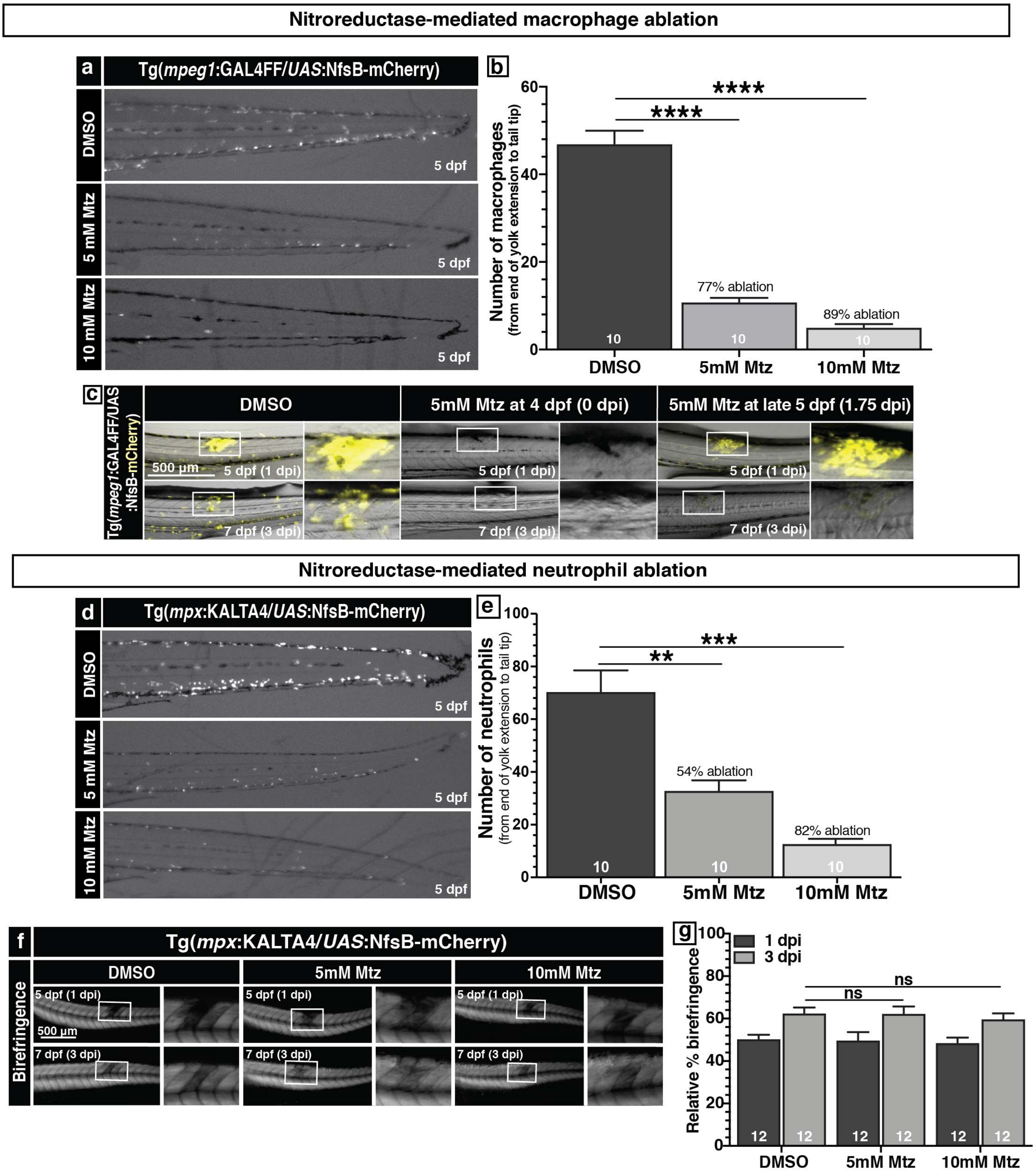

**Extended Data Figure 3**

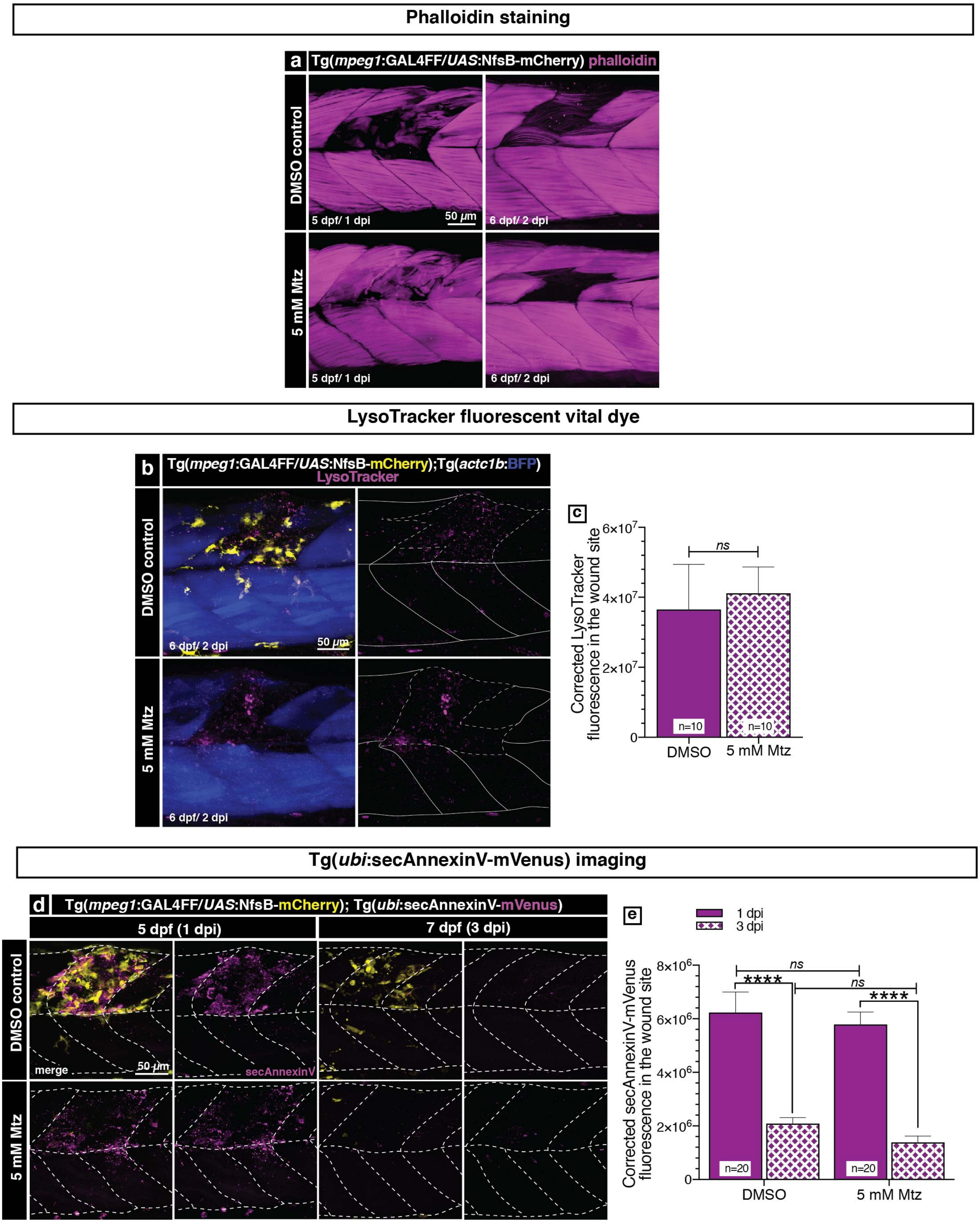

### Extended Data Figure 4

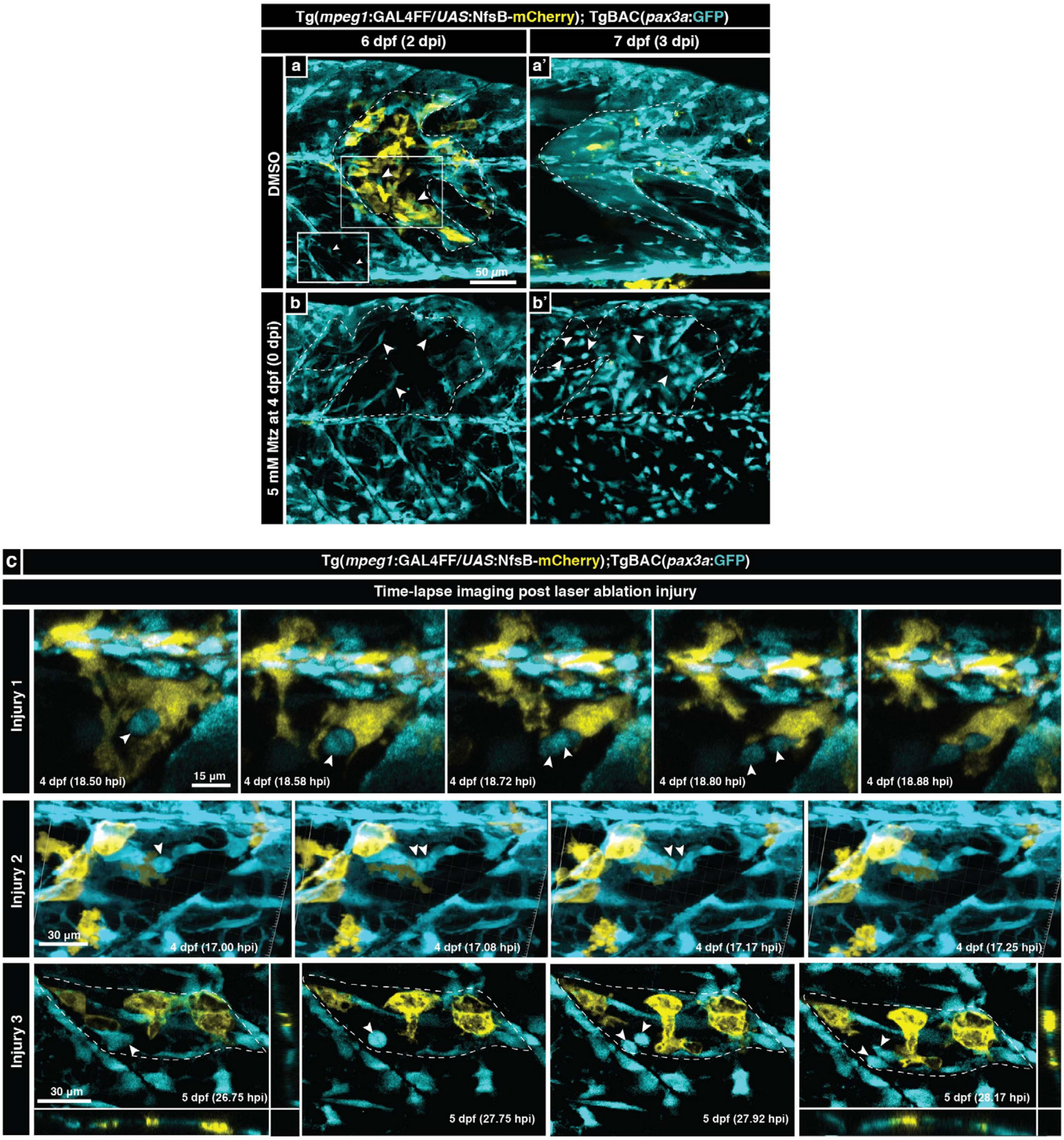

**Extended Data Figure 5**

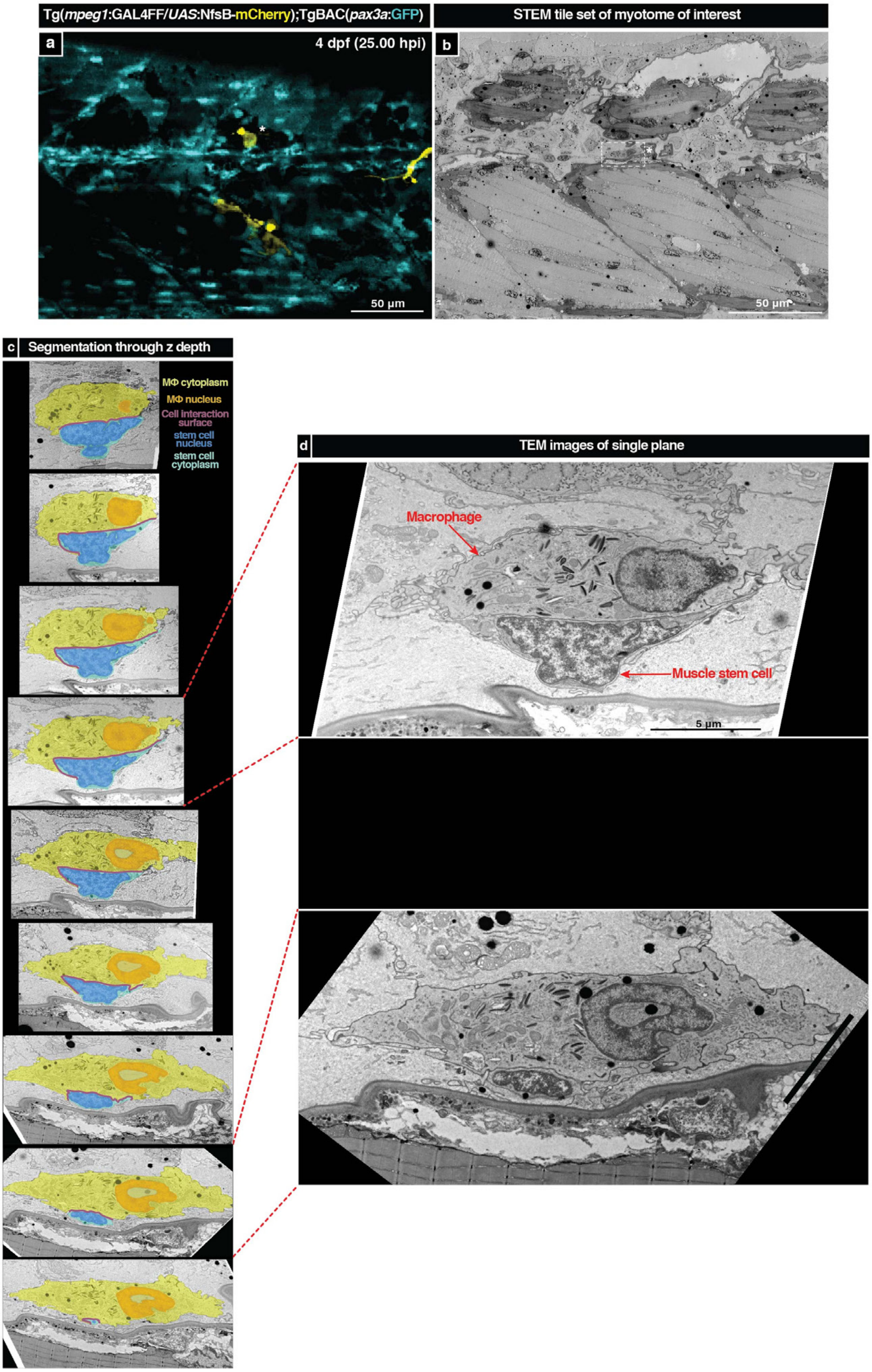

### Extended Data Figure 6

| # | Average logFC | p value | % cells in cluster 4 expressing gene | % cells outside cluster 4 expressing gene | Adjusted p value | Gene ID | Gene symbol |
|---|---|---|---|---|---|---|---|
| 1 | 3.51769726 | 2.46E-197 | 99.5% | 13.3% | 3.74E-193 | ENSDARG00000010423 | npsnl |
| 2 | 3.43652721 | 9.50E-143 | 100.0% | 33.8% | 1.44E-138 | ENSDARG00000042816 | mmp9 |
| 3 | 3.38223221 | 2.76E-185 | 98.6% | 14.8% | 4.18E-181 | ENSDARG00000019521 | mpx |
| 4 | 3.28014576 | 7.25E-150 | 98.6% | 25.7% | 1.10E-145 | ENSDARG00000012395 | mmp13a |
| 5 | 2.46606329 | 4.33E-208 | 91.1% | 6.0% | 6.57E-204 | ENSDARG00000041524 | NA |
| 6 | 2.16041847 | 6.24E-104 | 64.3% | 8.0% | 9.47E-100 | ENSDARG00000007769 | sult5a1 |
| 7 | 2.12152634 | 1.27E-168 | 82.2% | 7.0% | 1.94E-164 | ENSDARG00000007682 | ppdpfa |
| 8 | 2.10412796 | 1.31E-107 | 95.3% | 45.0% | 1.98E-103 | ENSDARG00000006029 | lta4h |
| 9 | 2.07830633 | 4.19E-152 | 90.6% | 14.2% | 6.36E-148 | ENSDARG00000090730 | cfbl |
| 10 | 1.87552844 | 3.06E-135 | 79.8% | 10.8% | 4.65E-131 | ENSDARG00000070398 | NA |
| 11 | 1.87151026 | 1.25E-87 | 76.5% | 19.9% | 1.91E-83 | ENSDARG00000089706 | si:ch211-276a23.5 |
| 12 | 1.80992908 | 1.77E-148 | 76.5% | 6.7% | 2.68E-144 | ENSDARG00000093124 | scpp8 |
| 13 | 1.7631956 | 1.24E-108 | 93.4% | 30.9% | 1.89E-104 | ENSDARG00000058348 | scinlb |
| 14 | 1.66944 | 3.32E-78 | 57.3% | 9.1% | 5.05E-74 | ENSDARG00000027852 | NA |
| 15 | 1.64420477 | 9.46E-67 | 68.1% | 19.4% | 1.44E-62 | ENSDARG00000033227 | lect2l |
| 16 | 1.63449553 | 1.20E-66 | 94.4% | 63.3% | 1.82E-62 | ENSDARG00000013598 | tnfb |
| 17 | 1.52901339 | 3.47E-94 | 96.2% | 56.7% | 5.28E-90 | ENSDARG00000035136 | selenow1 |
| 18 | 1.5127773 | 1.94E-97 | 83.1% | 21.4% | 2.95E-93 | ENSDARG00000078092 | limd2 |
| 19 | 1.51231421 | 2.59E-77 | 64.8% | 14.1% | 3.94E-73 | ENSDARG00000057430 | NA |
| 20 | 1.51190873 | 2.38E-123 | 66.2% | 5.9% | 3.62E-119 | ENSDARG00000056499 | ca6 |
| 21 | 1.50351826 | 1.06E-137 | 61.0% | 2.8% | 1.60E-133 | ENSDARG00000015657 | zgc:77112 |
| 22 | 1.49086617 | 7.47E-156 | 67.6% | 3.0% | 1.13E-151 | ENSDARG00000036382 | ponzr6 |
| 23 | 1.48791858 | 4.26E-93 | 97.2% | 60.3% | 6.46E-89 | ENSDARG00000077069 | srgn |
| 24 | 1.4694092 | 2.03E-114 | 100.0% | 99.4% | 3.08E-110 | ENSDARG00000095556 | LOC795545 |
| 25 | 1.45073072 | 1.23E-102 | 66.7% | 8.7% | 1.87E-98 | ENSDARG00000034187 | calm1b |
| 26 | 1.44369232 | 4.24E-67 | 53.5% | 9.3% | 6.44E-63 | ENSDARG00000039269 | arg2 |
| 27 | 1.4136587 | 1.17E-59 | 91.1% | 61.6% | 1.78E-55 | ENSDARG00000068784 | vsir |
| 28 | 1.37002123 | 7.17E-67 | 85.9% | 39.4% | 1.09E-62 | ENSDARG00000086337 | si:dkey-102g19.3 |
| 29 | 1.36152834 | 1.89E-57 | 60.1% | 15.2% | 2.87E-53 | ENSDARG00000078525 | noctb |
| 30 | 1.35387318 | 1.44E-69 | 88.3% | 42.5% | 2.18E-65 | ENSDARG00000007377 | odc1 |
| 31 | 1.34789643 | 5.30E-62 | 74.2% | 27.5% | 8.05E-58 | ENSDARG00000002021 | pygb |
| 32 | 1.33152195 | 4.44E-91 | 70.0% | 12.9% | 6.74E-87 | ENSDARG00000038458 | NA |
| 33 | 1.32613329 | 2.05E-120 | 52.6% | 2.1% | 3.12E-116 | ENSDARG00000086869 | NA |
| 34 | 1.30030857 | 4.46E-58 | 72.3% | 24.6% | 6.77E-54 | ENSDARG00000053836 | si:ch211-284o19.8 |
| 35 | 1.29743632 | 2.73E-71 | 89.2% | 45.1% | 4.15E-67 | ENSDARG00000063295 | myh9a |
| 36 | 1.26808592 | 1.71E-113 | 74.2% | 8.7% | 2.59E-109 | ENSDARG00000093193 | chia.6 |
| 37 | 1.25126448 | 6.86E-81 | 69.5% | 13.5% | 1.04E-76 | ENSDARG00000043457 | gapdh |
| 38 | 1.24510972 | 5.36E-77 | 64.8% | 13.2% | 8.14E-73 | ENSDARG00000052656 | si:ch211-193e13.5 |
| 39 | 1.23994196 | 8.64E-58 | 81.7% | 39.4% | 1.31E-53 | ENSDARG00000019236 | gsr |
| 40 | 1.22881477 | 1.20E-63 | 62.0% | 15.4% | 1.83E-59 | ENSDARG00000053381 | plpp1a |
| 41 | 1.22046143 | 2.53E-65 | 78.9% | 30.5% | 3.85E-61 | ENSDARG00000006019 | tktb |
| 42 | 1.22015324 | 9.85E-60 | 85.4% | 38.3% | 1.50E-55 | ENSDARG00000078547 | si:ch211-264f5.2 |
| 43 | 1.21978331 | 3.11E-80 | 100.0% | 71.9% | 4.72E-76 | ENSDARG00000031153 | NA |
| 44 | 1.21851595 | 5.37E-87 | 48.8% | 4.2% | 8.16E-83 | ENSDARG00000015922 | stk17al |
| 45 | 1.2094942 | 1.66E-99 | 59.6% | 6.3% | 2.52E-95 | ENSDARG00000009782 | myh11a |
| 46 | 1.19978784 | 2.62E-31 | 75.6% | 45.8% | 3.97E-27 | ENSDARG00000055186 | ccr9a |
| 47 | 1.19556383 | 1.95E-56 | 94.4% | 71.8% | 2.97E-52 | ENSDARG00000007693 | nfkbiab |
| 48 | 1.19263256 | 3.81E-64 | 49.3% | 7.9% | 5.78E-60 | ENSDARG00000013561 | pgm1 |
| 49 | 1.16112986 | 6.96E-69 | 93.0% | 56.2% | 1.06E-64 | ENSDARG00000052978 | mbnl1 |
| 50 | 1.16011422 | 1.77E-34 | 62.9% | 27.0% | 2.69E-30 | ENSDARG00000005673 | f3b |

| # | Average logFC | p value | % cells in cluster 4 expressing gene | % cells outside cluster 4 expressing gene | Adjusted p value | Gene ID | Gene symbol |
|---|---|---|---|---|---|---|---|
| 51 | 1.1314922 | 3.23E-53 | 85.4% | 49.3% | 4.90E-49 | ENSDARG00000014348 | stk17b |
| 52 | 1.12777964 | 7.32E-90 | 56.8% | 6.9% | 1.11E-85 | ENSDARG00000075870 | triobpa |
| 53 | 1.08027721 | 1.22E-59 | 57.7% | 12.8% | 1.86E-55 | ENSDARG00000032932 | cnksr1 |
| 54 | 1.07947549 | 6.74E-14 | 53.1% | 30.7% | 1.02E-09 | ENSDARG00000070622 | NA |
| 55 | 1.07860968 | 6.32E-37 | 48.8% | 13.9% | 9.60E-33 | ENSDARG00000018145 | mid1ip1l |
| 56 | 1.06525495 | 3.10E-70 | 61.5% | 12.0% | 4.71E-66 | ENSDARG00000059110 | NA |
| 57 | 1.05656185 | 3.64E-84 | 39.0% | 1.9% | 5.53E-80 | ENSDARG00000052766 | NA |
| 58 | 1.05024664 | 2.75E-45 | 77.5% | 42.4% | 4.18E-41 | ENSDARG00000026350 | wasb |
| 59 | 1.04344952 | 8.42E-38 | 78.9% | 45.7% | 1.28E-33 | ENSDARG00000035326 | nccrp1 |
| 60 | 1.02083795 | 3.73E-47 | 83.6% | 49.2% | 5.67E-43 | ENSDARG00000012987 | gpia |
| 61 | 1.01191026 | 6.86E-40 | 54.9% | 18.1% | 1.04E-35 | ENSDARG00000013705 | ccm2 |
| 62 | 1.00605145 | 8.59E-44 | 87.8% | 63.9% | 1.30E-39 | ENSDARG00000036967 | smox |
| 63 | 1.00599877 | 4.72E-50 | 59.6% | 16.4% | 7.17E-46 | ENSDARG00000070601 | si:dkey-85p17.3 |
| 64 | 0.99619739 | 4.52E-40 | 59.6% | 22.4% | 6.86E-36 | ENSDARG00000071065 | g6pd |
| 65 | 0.9934688 | 1.82E-30 | 83.6% | 56.3% | 2.76E-26 | ENSDARG00000078619 | pnp5a |
| 66 | 0.99273801 | 3.21E-43 | 82.6% | 53.1% | 4.88E-39 | ENSDARG00000012256 | NA |
| 67 | 0.97506782 | 5.09E-33 | 79.8% | 53.2% | 7.73E-29 | ENSDARG00000044619 | birc2 |
| 68 | 0.97451848 | 8.87E-95 | 100.0% | 100.0% | 1.35E-90 | ENSDARG00000037870 | actb2 |
| 69 | 0.97310907 | 9.40E-39 | 51.6% | 15.1% | 1.43E-34 | ENSDARG00000001452 | adam8a |
| 70 | 0.97102903 | 3.35E-48 | 90.1% | 59.4% | 5.08E-44 | ENSDARG00000062721 | dennd4a |
| 71 | 0.96769076 | 4.35E-32 | 66.2% | 32.4% | 6.61E-28 | ENSDARG00000030598 | nampta |
| 72 | 0.96659516 | 1.88E-80 | 41.8% | 2.8% | 2.85E-76 | ENSDARG00000055705 | f5 |
| 73 | 0.96522809 | 5.73E-92 | 50.2% | 4.0% | 8.69E-88 | ENSDARG00000060149 | ablim1a |
| 74 | 0.96018943 | 1.02E-84 | 39.4% | 1.8% | 1.55E-80 | ENSDARG00000077975 | si:ch73-343l4.8 |
| 75 | 0.95761111 | 1.31E-23 | 62.4% | 34.7% | 1.99E-19 | ENSDARG00000042382 | NA |
| 76 | 0.95416288 | 9.99E-37 | 76.1% | 41.5% | 1.52E-32 | ENSDARG00000036637 | arl11 |
| 77 | 0.95084342 | 3.31E-36 | 47.4% | 14.0% | 5.03E-32 | ENSDARG00000079353 | nmrk1 |
| 78 | 0.94000322 | 5.85E-34 | 81.2% | 62.8% | 8.88E-30 | ENSDARG00000068214 | ccni |
| 79 | 0.93656916 | 1.41E-42 | 77.5% | 36.2% | 2.14E-38 | ENSDARG00000004979 | elovl5 |
| 80 | 0.93194757 | 1.38E-28 | 87.8% | 64.3% | 2.09E-24 | ENSDARG00000033735 | ncf1 |
| 81 | 0.92953484 | 1.09E-41 | 55.9% | 17.8% | 1.66E-37 | ENSDARG00000057053 | zgc:91976 |
| 82 | 0.92892269 | 8.26E-57 | 46.0% | 7.4% | 1.25E-52 | ENSDARG00000010556 | mmp25b |
| 83 | 0.92427477 | 8.88E-71 | 58.7% | 10.9% | 1.35E-66 | ENSDARG00000004647 | spice1 |
| 84 | 0.92319626 | 3.75E-40 | 54.5% | 17.1% | 5.70E-36 | ENSDARG00000063195 | pxk |
| 85 | 0.91734144 | 6.07E-37 | 65.7% | 26.6% | 9.22E-33 | ENSDARG00000039884 | NA |
| 86 | 0.91012995 | 1.49E-29 | 33.8% | 7.9% | 2.26E-25 | ENSDARG00000012671 | inhbaa |
| 87 | 0.9082643 | 2.54E-102 | 43.7% | 1.5% | 3.85E-98 | ENSDARG00000069440 | dachd |
| 88 | 0.9067071 | 5.33E-64 | 37.1% | 3.1% | 8.09E-60 | ENSDARG00000009208 | prkcda |
| 89 | 0.90038508 | 1.59E-25 | 66.7% | 36.9% | 2.41E-21 | ENSDARG00000019702 | aldocb |
| 90 | 0.89617091 | 2.31E-103 | 43.2% | 1.4% | 3.51E-99 | ENSDARG00000093248 | NA |
| 91 | 0.89200683 | 6.01E-32 | 77.9% | 48.4% | 9.12E-28 | ENSDARG00000054272 | caprin1b |
| 92 | 0.8789271 | 1.17E-55 | 42.3% | 6.3% | 1.78E-51 | ENSDARG00000043182 | hhla2a.1 |
| 93 | 0.87828949 | 9.16E-54 | 95.8% | 83.4% | 1.39E-49 | ENSDARG00000027063 | arpc1b |
| 94 | 0.87679857 | 3.35E-30 | 68.1% | 35.8% | 5.08E-26 | ENSDARG00000010752 | acsl4b |
| 95 | 0.87632591 | 5.80E-34 | 68.1% | 30.9% | 8.80E-30 | ENSDARG00000071384 | slc7a11 |
| 96 | 0.86981487 | 5.22E-40 | 68.5% | 30.1% | 7.93E-36 | ENSDARG00000004378 | yrk |
| 97 | 0.86948795 | 6.80E-63 | 100.0% | 95.1% | 1.03E-58 | ENSDARG00000088091 | pfn1 |
| 98 | 0.86896945 | 1.20E-37 | 32.4% | 5.5% | 1.82E-33 | ENSDARG00000053931 | si:dkey-42l23.5 |
| 99 | 0.86896495 | 3.74E-48 | 90.1% | 60.8% | 5.68E-44 | ENSDARG00000041505 | itm2bb |
| 100 | 0.86866486 | 1.59E-16 | 66.2% | 44.8% | 2.41E-12 | ENSDARG00000056875 | NA |

**Extended Data Figure 7**

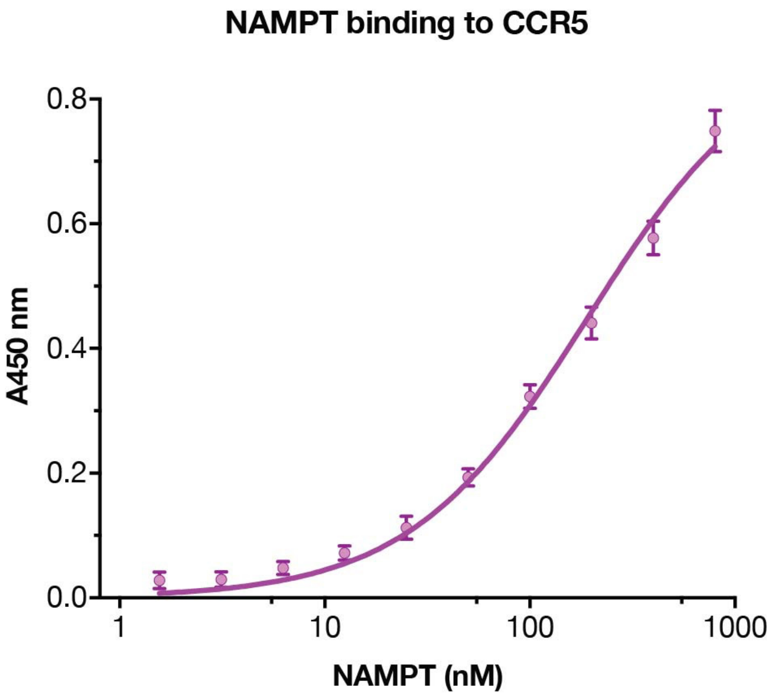

**Extended Data Figure 8**

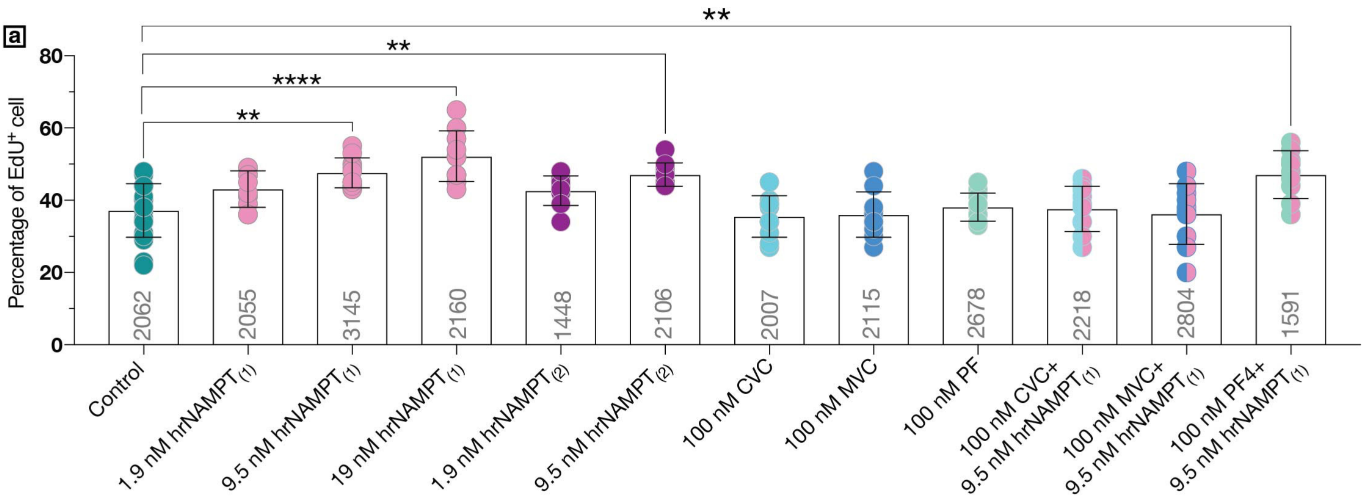

### Extended Data Figure 9

a

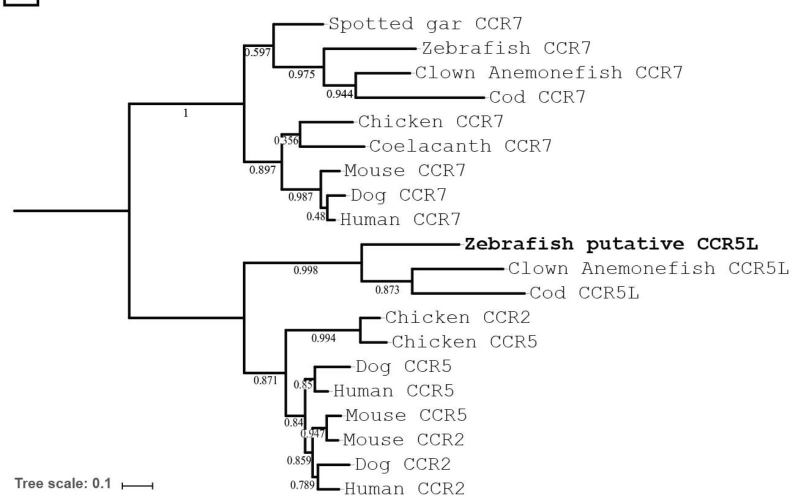


| Species/protein | Accession Number |
|---|---|
| Spotted gar CCR7 | ENSLOCT00000022115.1 |
| Zebrafish CCR7 | NP_001092213.1 |
| Clown Anemonefish CCR7 | ENSAOCT00000017280.1 |
| Cod CCR7 | ENSGMOT00000022477.1 |
| Chicken CCR7 | NP_001185681.1 |
| Coelacanth CCR7 | ENSLACT00000015269.2 |
| Mouse CCR7 | NP_001288642.1 |
| Dog CCR7 | ENSCAFT00000049953.1 |
| Human CCR7 | NP_001829.1 |
| Zebrafish putative CCR5L | XP_001344151.1 |
| Clown Anemonefish CCR5L | ENSAOCG00000012993 |
| Cod CCR5L | ENSGMOT00000005070.1 |
| Chicken CCR2 | NP_001039300.1 |
| Chicken CCR5 | NP_001258070.1 |
| Dog CCR5 | NP_001012342.2 |
| Human CCR5 | NP_000570.1 |
| Mouse CCR5 | NP_034047.2 |
| Mouse CCR2 | NP_034045.1 |
| Dog CCR2 | E2QTN7 |
| Human CCR2 | NP_001116513.2 |

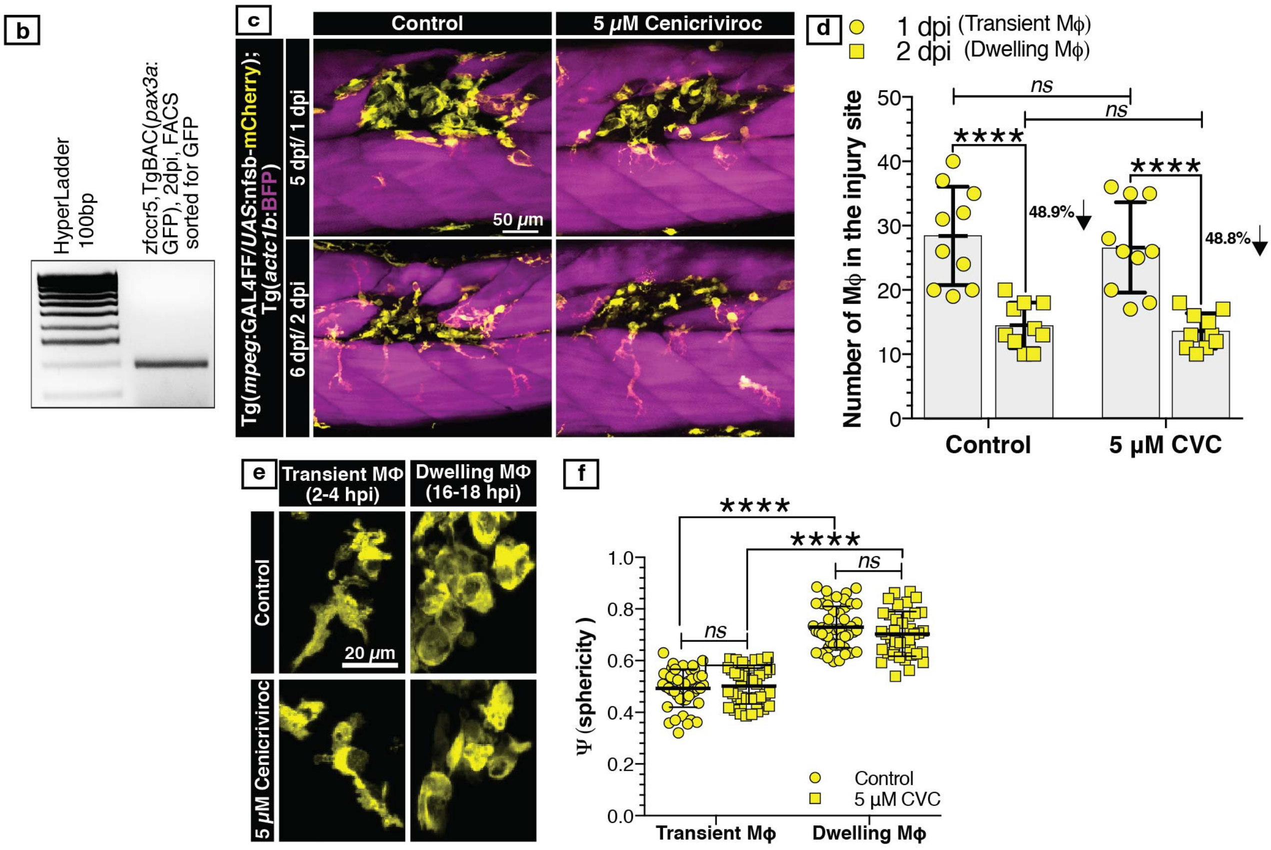

### Extended Data Figure 10

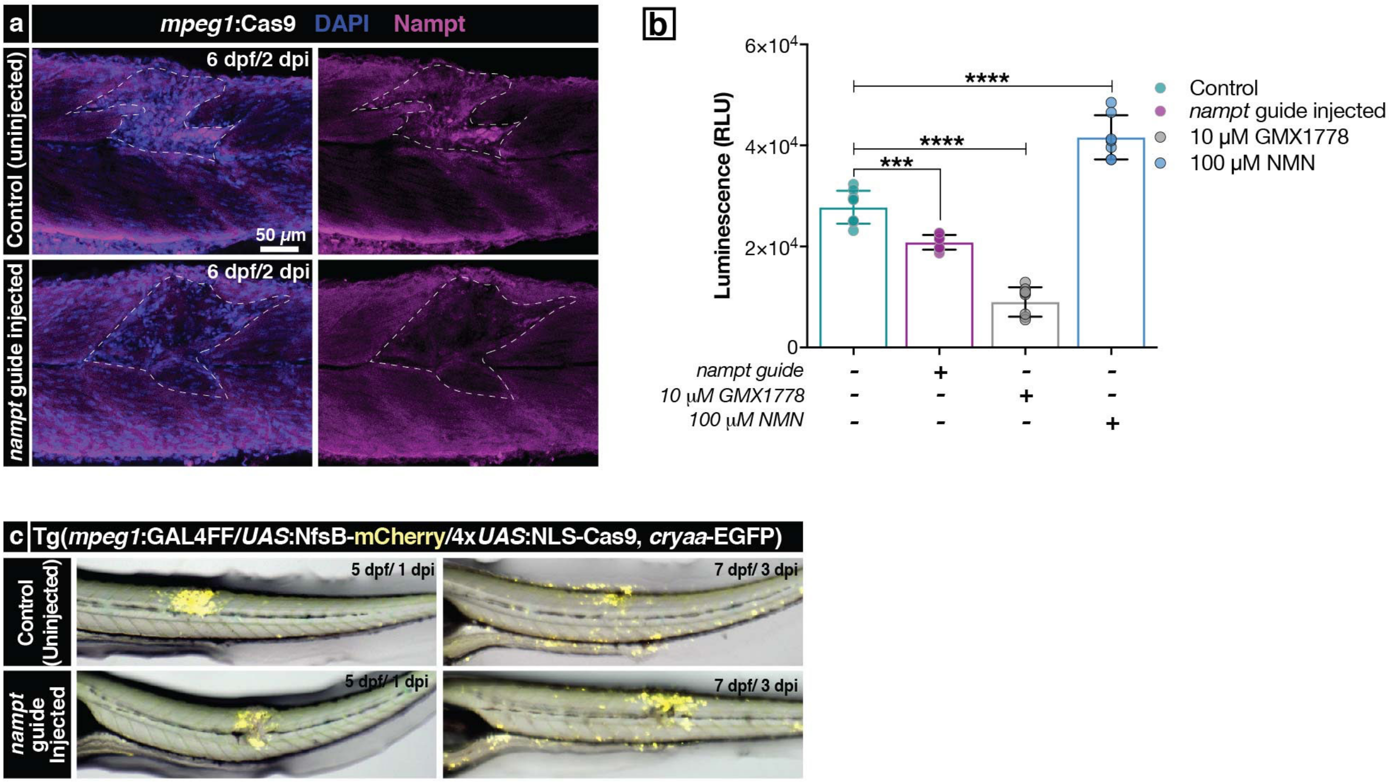
